# BioMeBar Indian Gut Reference Atlas: A Harmonised Multi-Cohort Healthy Microbiome Baseline and the Indian Gut Health Index

**DOI:** 10.64898/2026.10.11.758353

**Authors:** Tushar Parihar, Nikita Dinger, Rahul Makhijani

## Abstract

Population-scale gut microbiome reference databases exist for Western and East Asian cohorts, yet no harmonised healthy baseline has been available for India’s 1.4 billion inhabitants, whose diets span extreme geographic and cultural gradients. Western-derived reference ranges risk systematic misclassification when applied to Indian subjects.

We harmonised eight published Indian healthy-gut microbiome cohorts (1,672 samples from 1,456 unique individuals, 20 geographic locations, six zones) onto a common NCBI family-level taxonomy axis. Raw 16S rRNA amplicon data were reprocessed through a uniform QIIME 2 / DADA2 / SILVA 138.1 pipeline where raw sequences were available; author-provided taxonomic tables from MetaPhlAn, HITChip, and RDP-based pipelines were retained and mapped via a four-step resolution cascade (exact match, synonym lookup, genus-to-family fallback, Unassigned). We defined the Indian Gut Health Index (IGHI), a four-component composite score (diversity, taxon balance, core membership, community typicality) calibrated against geography-stratified healthy Indian norms. The unified matrix comprised 380 bacterial families, of which 17 constituted the pan-Indian core (≥80% prevalence; *Lachnospiraceae* 100%, *Prevotellaceae* 99.5%). PERMANOVA confirmed significant compositional differences across geographies (pseudo-F = 43.4, p = 0.001) and cohorts (pseudo-F = 89.6, p = 0.001). Variance partitioning revealed that geography and cohort effects were deeply confounded (shared variance 30.4%; pure geography 4.1%, pure cohort 4.7%, residual 60.9%). Restricted permutation PERMANOVA confirmed city-level geographic signal after cohort correction (p = 0.001), but zone-level signal did not survive (p = 0.896). Two community types were identified: a *Prevotellaceae*-dominant type (69%) and an *Oscillospiraceae*-dominant type (31%), contrasting with the Western three-enterotype model. A Random Forest zone classifier achieved 68.4% balanced accuracy (null 16.6%), with Northeast subjects most distinguishable (F1 = 0.92). IGHI zone means ranged from 67.6 to 72.9 (inter-zone range 5.3 points), confirming effective geographic normalisation. Weight sensitivity analysis showed IGHI rankings were robust to moderate perturbation (Spearman ρ ≥ 0.88 for five of six alternative schemes).

The BioMeBar^TM^ Indian Gut Reference Atlas provides the first pan-Indian harmonised healthy microbiome baseline, and the IGHI offers a population-calibrated scoring framework for gut health assessment in Indian individuals. The atlas, scoring model, and analysis code are openly available at https://github.com/TusharParihar128/BioMeBar_Indian_Healthy_Cohorts_datasets.

## 1. Introduction

The human gut microbiome is now recognised as a critical determinant of metabolic, immunological, and neurological health. Large-scale reference projects - including the Human Microbiome Project (HMP; [1], the MetaHIT consortium [2], the Flemish Gut Flora Project [3], and the American Gut Project [4] have established population baselines that anchor clinical interpretation and discovery in their respective cohorts. More recently, the Unified Human Gastrointestinal Genome [5] and the Human Reference Gut Microbiome [6] have extended reference coverage to thousands of metagenome-assembled genomes. Yet the overwhelming majority of subjects contributing to these resources are of European or East Asian ancestry. India, home to one-sixth of the world’s population, remains conspicuously under-represented. In fact, Indians represent less than 1 to 1.8% of samples in global human microbome studies [7].

This gap is not merely academic. Gut microbiome composition in Indian populations differs substantially from Western references along multiple axes. Prevotella-dominant enterotypes, associated with high-fibre plant-based diets, are markedly more prevalent in Indian cohorts than in industrialised Western populations where Bacteroides-dominant profiles predominate [8], [9]. The Indian diet spans an extraordinary range, from the strictly lacto-vegetarian Jain communities of Gujarat to the fermented-fish diets of the Meitei people in Manipur, and this dietary diversity is also reflected in microbial variation in the gut that cannot be captured by any Western-calibrated scoring systems. Commercially available gut health tests in India currently report against such Western reference distributions. They hence risk systematic misclassification when applied to Indians: a Prevotella-rich profile that falls within the normal Indian range may be flagged as aberrant against a Bacteroides-centric Western baseline.

Several independent studies have characterised healthy Indian gut microbiomes, but each has been limited to a single geographic region, a single sequencing platform, or a single taxonomic resolution. Dhakan et al. (2019)[8] profiled 110 subjects from Bhopal and Kerala using both 16S amplicon and whole-genome shotgun sequencing, reporting diet-associated compositional differences and a fecal plus serum metabolomics layer. Dubey et al. (2018)[10] established the LogMPIE pan-India gut microbiome resource by profiling 1,004 healthy individuals across 14 geographic locations in India using Ion S5-based 16S rRNA sequencing. Tandon et al. (2018)[11] characterised the gut microbiome of 80 healthy urban individuals from Ahmedabad, with paired stool samples collected at two time points approximately 60 days apart, using RDP-based taxonomic classification and Vikodak for functional profiling. Bhute et al. (2016)[12] characterised gut microbial communities in 34 healthy Indian subjects from Delhi and Pune and nearby rural regions, reporting enrichment of Prevotella and Megasphaera. Kumbhare et al. (2017)[13] compared 47 Indian and 52 Finnish adolescents from Pune, with PICRUSt-inferred functional predictions. Chaudhari et al. (2020)[14] characterised the gut, oral and skin microbiomes of 54 healthy members of six three-generation patrilineal families from the Pune region using 16S rRNA sequencing; after quality-based exclusion, 50 gut samples were retained for analysis. Kaur et al. (2020)[9] provided MetaPhlAn-classified multi-city profiles with detailed dietary metadata. Jeyaram et al. (2025)[15] characterised 78 agrarian Meitei individuals from Manipur using HITChip phylogenetic microarray and untargeted LC-HRMS fecal metabolomics, reporting 871 detected metabolite features. D’Aloisio et al. (2025)[16] profiled 110 subjects spanning India-resident Indians, Indo-Immigrants, and Indo-Canadians using both 16S rRNA and whole-genome shotgun sequencing, providing the atlas’s cross-platform validation cohort.

No prior work has harmonised these cohorts into a single reference frame. The studies differ in sequencing platform (Illumina MiSeq, Ion Torrent PGM, HITChip microarray), bioinformatics pipeline (QIIME 1.x, QIIME 2/DADA2, MetaPhlAn, RDP Classifier), reference database (GreenGenes, SILVA, RDP), and taxonomic resolution (OTU vs. ASV vs. genus-level vs. probe-based groups). Direct comparison across cohorts is therefore impossible without a deliberate harmonisation strategy.

Here we present the BioMeBar^TM^ Indian Gut Reference Atlas: a harmonised, family-level taxonomic matrix spanning eight cohorts, approximately 1,672 samples from 1,456 unique individuals, and 20 geographic locations across India. We adopt NCBI family-level taxonomy as the common resolution - the coarsest level that preserves biologically meaningful distinctions while absorbing the heterogeneity introduced by different classifiers and reference databases. We define and compute the Indian Gut Health Index (IGHI), a composite scoring model calibrated against geography-stratified healthy Indian norms, designed for clinical and commercial gut health assessment in Indian populations. The atlas, scoring model, and analysis code are openly available at https://github.com/TusharParihar128/BioMeBar_Indian_Healthy_Cohorts_datasets.

## 2. Methods

### 2.1 Literature identification and cohort selection

We conducted a structured literature search across PubMed, Google Scholar, and NCBI BioProject to identify published studies characterising the gut microbiome of healthy Indian adults. Search terms included combinations of “gut microbiome”, “gut microbiota”, “16S rRNA”, “metagenome”, “India”, “Indian”, and specific geographic regions. Studies were included if they (i) profiled the fecal microbiome of healthy Indian subjects (no diagnosed gastrointestinal disease at time of sampling), (ii) provided taxonomic abundance data at genus level or higher, and (iii) made raw sequence data or processed abundance tables publicly available.

Eight cohorts met inclusion criteria (Table 1). Studies were categorised into two processing tracks: those requiring *de novo* sequence processing from raw SRA deposits, and those for which author-provided taxonomic tables could be used directly after harmonisation onto the project-wide taxonomy axis.

**Table 1.** Included cohorts.

| Cohort | Reference | n | Geography | Sequencing | Classifier | Data source |
| --- | --- | --- | --- | --- | --- | --- |
| Dhakan | Dhakan et al., 2019[8] | 110 | Bhopal, Kerala | 16S + WGS | SILVA 132 / Bowtie-2 | PRJNA397112, GigaDB |
| LogMPIE | Dubey et al., 2018[10] | 1,004 | Pan-India (14 locations) | 16S (Ion Torrent) | MEGABLAST + Greengenes | PRJEB25642 (ENA) |
| Kamaldeep | Kaur et al., 2020[9] | 31 | Ladakh, Jaisalmer & Khargone | WGS (shotgun metagenomics) | MetaPhlAn2 / OneCodex | PRJNA531203 |
| Tandon | Tandon et al., 2018[11] | 80 | Ahmedabad | 16S | RDP Classifier v2.11 | Author tables |
| Jeyaram | Jeyaram et al., 2025[15] | 78 | Manipur | HITChip microarray | HITChip probe-based | Zenodo 10.5281/zenodo.14424024 |
| Kumbhare | Kumbhare et al., 2017[13] | 47 (Indian) | Pune | 16S | GreenGenes 13.8 / RDP | SRP077882/ PRJNA327770 |
| Chaudhari | Chaudhari et al., 2020[14] | 54 | Pune | 16S | GreenGenes 13.8 | SRP116277/ PRJNA399246 |
| D'Aloisio | D'Aloisio et al., 2025[16] | 61 India n + 32 Indo-Immigrant + 17 Indo-Canadian (110 total) | India-based Indians, Indo-Immigrants, and Indo-Canadians in Canada | 16S & WGS | GreenGenes 2 & MetaPhlAn4 | PRJNA1082632 |

### 2.2 Raw sequence acquisition and processing

For the Kumbhare, Chaudhari, and D’Aloisio cohorts, publicly available raw 16S rRNA sequencing reads were retrieved and reprocessed through a reproducible QIIME 2–DADA2 workflow. Raw-read quality was assessed with FastQC and summarised using MultiQC, followed by dataset-specific preprocessing and denoising parameters selected according to the sequencing-read structure and observed quality profiles. Paired-end reads were processed using the QIIME 2 DADA2 plugin for sequence denoising, paired-read merging, chimera removal, and ASV inference, with trimming and truncation settings applied separately for each cohort. Taxonomic classification was subsequently performed using the QIIME 2 classify-sklearn Naive Bayes framework with a SILVA 138.1 SSURef_NR99 reference classifier **[17], [18], [19].**

**Table 2.** Cohort Processing Workflows.

| Cohort | Raw 16S data processed | FastQC | MultiQC | QIIME 2 / DADA2 | Dataset-specific preprocessing / DADA2 settings | Taxonomic classification |
| --- | --- | --- | --- | --- | --- | --- |
| Kumbhare et al., 2017[13] | 42 paired-end runs | v0.11.9 | v1.35 | QIIME 2 release 2024.10; q2cli 2024.10.1; DADA2 plugin 2024.10.0 | Cutadapt primer removal; DADA2 **denoise-paired**; trim-left 0/0 after primer removal; truncation 200/160 bp; max EE 2/2; truncation quality 2; minimum overlap 12 bp; independent pooling; consensus chimera removal | SILVA 138.1 / SSURef_NR99 Naive Bayes, confidence 0.7 |
| Chaudhari et al., 2020[14] | 50 paired-end gut runs | v0.11.9 | v1.35 | QIIME 2 release 2024.10; q2cli 2024.10.1; DADA2 paired-end workflow | Final Cutadapt trimming on R1-derived sequences; minimum length 250 nt; DADA2 **denoise-single**; **trunc_len=0**; max EE 2.0; trim-left 0; consensus chimera removal; no explicit read-merging command in execution history | SILVA 138.1 / SSURef_NR99 Naive Bayes, **classify-sklearn**, confidence 0.7 |
| D'Aloisio et al., 2025[16] | 110 paired-end 16S runs | v0.12.1 | v1.35 | QIIME 2 release 2024.10; q2cli 2024.10.1; DADA2 | DADA2 **denoise-paired**; trim-left 31/32 bp; truncation 129/205 bp; **n_threads=0**; no | SILVA 138.1 / SSURef_NR99 Naive Bayes, **classify-sklearn**, n_jobs=8 |
|  |  |  |  | paired-end workflow | separate Cutadapt step documented |  |

For the Kamaldeep cohort (Kaur et al., 2020)[9], author-provided MetaPhlAn2-classified family-level abundance tables were used directly, as the study employed both 16S and whole-genome sequencing with a clade-specific marker gene approach not reproducible from 16S reads alone.

For the Jeyaram cohort (Jeyaram et al., 2025)[15], HITChip phylogenetic microarray probe-level data were obtained from Zenodo. HITChip reports taxonomy as “level-1” (order-like, >90% 16S sequence similarity groups) and “level-2” (genus-like groups). Level-2 genus-like groups were mapped to NCBI families via the genus-to-family lookup described in Section 2.4.

For the Tandon cohort (Tandon et al., 2018)[11], RDP Classifier v2.11-assigned taxonomy at 80% bootstrap confidence was used as provided by the authors.

The original studies employed different taxonomic classifiers: Kumbhare et al. (2017)[13] used GreenGenes 13.8 with the RDP Classifier, Chaudhari et al. (2020)[14] used GreenGenes 13.8, and D’Aloisio et al. (2025)[16] used GreenGenes2. However, as processed taxonomic tables from these original analyses were not publicly available, we reprocessed all raw sequences through our standardised QIIME 2 / DADA2 / SILVA 138.1 NR99 Naive Bayes workflow to ensure consistent taxonomic assignment across all cohorts included in the atlas. Table 1 reports our classifier (SILVA 138.1 Naive Bayes) for these three cohorts, reflecting the taxonomy actually used in the harmonised matrix.

### 2.3 Functional prediction

For the Kumbhare, Chaudhari, and D’Aloisio cohorts, functional potential was inferred from de novo QIIME 2-generated ASV feature tables using PICRUSt2, with **v2.6.3 for Kumbhare and Chaudhari and v2.5.2 for D’Aloisio** [20]. The pipeline comprised:

1. **Phylogenetic placement** of study ASVs into the PICRUSt2 reference tree [21].
2. **Hidden-state prediction** of gene family copy numbers per ASV (castor R package integration).
3. **Metagenome prediction** by weighting predicted gene copies by ASV abundance.
4. **Pathway inference** using MinPath [22] to predict MetaCyc pathway abundances.

Output tables included EC number abundances, KEGG Ortholog (KO) abundances, and MetaCyc pathway abundances per sample. Nearest Sequenced Taxon Index (NSTI) values were computed for each ASV; samples with mean weighted NSTI > 0.15 were flagged for cautious interpretation.

The Dhakan cohort’s functional profile was derived from the original study’s whole-genome shotgun analysis (Bowtie-2 mapping against 4,097 HMP/NCBI reference genomes, with eggNOG and KEGG Ortholog annotation via the Indian Microbial Gene Catalogue). The Kamaldeep cohort’s functional layer comprised CAZyme profiles from *de novo* assembly, MetaGeneMark gene prediction, and dbCAN/HMMscan classification. These study-specific functional analyses were retained as provided, without re-processing.

### 2.4 NCBI family-level taxonomic harmonisation

The central challenge of this work was projecting taxonomic labels from seven different classification systems onto a single axis. We selected NCBI Taxonomy as the reference backbone and family rank as the resolution target, for three reasons:

1. **Family rank absorbs classifier disagreement.** Two classifiers may assign the same ASV to different genera within the same family (e.g., *Roseburia* vs. *Eubacterium* within Lachnospiraceae); family-level aggregation resolves this without information loss at the biologically interpretable level.
2. **Family rank spans all input platforms.** HITChip level-2 genus-like groups, MetaPhlAn clade-specific markers, and OTU/ASV-based classifiers all resolve reliably to family, whereas species or even genus resolution is inconsistent across platforms.
3. **NCBI Taxonomy is the most complete backbone.** GreenGenes 13.8 (last updated 2013) and SILVA use their own taxonomic frameworks; NCBI provides a continuously updated, comprehensive mapping that subsumes both.

The harmonisation cascade operated as follows, applied to every unique taxonomic label in every cohort:

1. **Exact match.** The label was looked up directly in NCBI Taxonomy at family rank.
2. **Synonym resolution.** If no exact match, the label was checked against the NCBI synonym table (including historical names, misspellings, and reclassified taxa).
3. **Genus-to-family fallback.** If the label was a genus name (common for MetaPhlAn and HITChip outputs), the genus was looked up in NCBI and its parent family extracted.
4. **Unassigned.** Labels that could not be resolved at any step were flagged as “Unassigned” and retained in the matrix to preserve compositional closure.

This cascade was implemented programmatically with manual verification of ambiguous cases. The full mapping table-input label, matched NCBI family, resolution step used, and any manual overrides is deposited with the dataset.

After harmonisation, per-sample abundance vectors were re-normalised to sum to 1.0 (compositional closure) to account for the redistribution of abundance from split or merged taxa.

### 2.5 Unified matrix construction

The harmonised family-level abundance tables from all cohorts were concatenated into a single matrix (subjects × families). Each row carries four metadata columns: cohort (study identifier), geography (geographic region), sequencing (platform type: 16S-Illumina, 16S-IonTorrent, HITChip, WGS), and timepoint (for the longitudinal Tandon and Jeyaram cohorts; baseline for all others).

Two cohorts contributed longitudinal samples: Tandon (80 subjects × 2 timepoints, 60 days apart) and Jeyaram (78 subjects × up to 3 seasonal timepoints), yielding 1,672 total samples from 1,456 unique individuals. Statistical tests that assume sample independence are reported on the full dataset; a sensitivity analysis using one randomly selected sample per individual confirmed robustness of all primary findings.

Two versions of the matrix were produced:

- **Union matrix:** all families observed in any cohort (sparse; zeroes indicate absence or below-detection).
- **Common matrix:** intersection of families observed in every cohort (dense; useful for cross-cohort comparisons without zero-inflation artefacts).

The matrix auto-detection routine handles both orientations (subjects × families and families × subjects), transposing if necessary, and verifies compositional closure (row sums within 1 ± 0.01 after normalisation).

### 2.6 Indian Gut Health Index (IGHI) definition

We define the IGHI as a weighted composite of four components, each scored on a 0–100 scale:

#### Component 1: Diversity Score (weight = 0.30)

The Shannon diversity index [23] is computed from the family-level abundance vector for each subject. The raw Shannon value is converted to a percentile rank *within the subject’s geographic stratum*, yielding a score of 0–100 where 100 indicates the highest diversity observed among healthy individuals from the same region. Geography-stratified percentiling accounts for the known influence of geography on baseline diversity - a Shannon index that is normal for Manipur may be low for Pune - preventing geographic confounding from biasing the score.

#### Component 2: Taxon Balance Score (weight = 0.25)

This component evaluates the ratio of beneficial to potentially opportunistic families in an individual’s gut profile. Beneficial families were selected based on consistent positive association with health outcomes in Indian and global literature: Lachnospiraceae, Ruminococcaceae, Bifidobacteriaceae, Lactobacillaceae, Prevotellaceae, and Erysipelotrichaceae. Opportunistic families - those enriched in dysbiotic states - include Enterobacteriaceae, Staphylococcaceae, Pseudomonadaceae, and Clostridiaceae. The score is computed as:

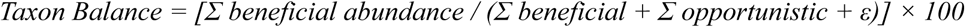

where ε = 10⁻⁶ prevents division by zero. This ratio is robust to differences in absolute abundance across sequencing depths and normalisation approaches.

#### Component 3: Core Membership Score (weight = 0.20)

Core Indian gut families are defined as those present (relative abundance > 0) in ≥80% of all subjects in the atlas. The core membership score for an individual is the fraction of core families detected in their sample, scaled to 0–100. A subject harbouring all core families scores 100; one missing half of them scores 50. If no family meets the 80% prevalence threshold (unlikely given the atlas size), the threshold relaxes to 50%.

#### Component 4: Community Typicality Score (weight = 0.25)

For each geographic stratum, a centroid is computed as the mean family-abundance vector across all healthy subjects from that region. Each subject’s Bray-Curtis dissimilarity (Bray & Curtis, 1957)[24] to their stratum centroid is calculated and inverted:

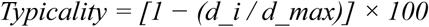

where *d_i* is the subject’s distance to centroid and *d_max* is the maximum distance observed in that stratum. Subjects closest to the healthy centroid score highest; outliers score lowest.

#### Composite IGHI

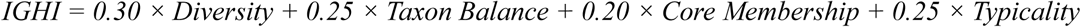

Default weights were assigned based on the relative informativeness of each component in preliminary analysis and literature precedent: diversity is the most widely validated single-metric correlate of gut health, hence the highest weight; taxon balance and community typicality capture complementary dimensions of compositional health; core membership provides a simple presence/absence check against the Indian-specific reference. Weights are user-configurable in the open-source implementation to support sensitivity analysis and future validation studies.

**Interpretive thresholds** (preliminary, pending validation against disease cohorts):

● 80–100: Excellent gut health profile (top quintile of Indian reference)

● 60–79: Good - within healthy Indian range

● 40–59: Moderate - some deviation from healthy reference

● 20–39: Below average - significant deviation

● 0–19: Low - consider clinical evaluation

These thresholds are derived from the distribution of IGHI scores in the healthy atlas and should be validated against Indian IBS and other gastrointestinal disease cohorts before clinical deployment.

### 2.7 Statistical analysis

All statistical analyses were performed in Python 3.11 using pandas (v2.0+), NumPy (v1.24+), SciPy (v1.10+), scikit-bio (v0.5.9+), scikit-learn (v1.3+), and statsmodels (v0.14+).

#### Alpha diversity

Shannon diversity, Simpson’s diversity, observed family richness, and Pielou’s evenness were computed per sample from the unified family matrix. Differences across geographies and cohorts were tested using the Kruskal-Wallis H test (non-parametric, appropriate for non-normally distributed ecological data), with post-hoc pairwise Mann-Whitney U tests and Benjamini-Hochberg false discovery rate (FDR) correction at q < 0.05.

#### Beta diversity

Pairwise Bray-Curtis dissimilarity was computed across all subjects. Principal Coordinates Analysis (PCoA) was performed on the distance matrix using scikit-bio, and ordination results were visualised as 2D scatter plots coloured by geography. The contribution of geography and cohort identity to compositional variation was assessed using Permutational Multivariate Analysis of Variance [25], implemented via skbio.stats.distance.permanova with 999 permutations.

#### Differential abundance

Families differentially abundant across geographic regions were identified using the Kruskal-Wallis test applied to each family independently, with Benjamini-Hochberg FDR correction. For significant families (q < 0.05), pairwise Mann-Whitney U tests were performed between all geography pairs with n ≥ 5 subjects per group. Effect size was quantified using Cliff’s delta [26], a non-parametric measure bounded between −1 and +1 that is robust to non-normality and unequal sample sizes:

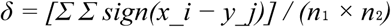

where |δ| < 0.147 is negligible, 0.147–0.33 is small, 0.33–0.474 is medium, and > 0.474 is large.

#### Geography classification

A Random Forest classifier [27] with 500 trees, balanced class weights (to address unequal group sizes), and minimum 3 samples per leaf was trained to predict geographic origin from family-level abundance profiles. Model performance was assessed using stratified 5-fold cross-validation (accuracy and balanced accuracy) and leave-one-cohort-out cross-validation (to assess generalisability beyond batch effects). Feature importances (Gini impurity decrease) were extracted to identify the most geography-discriminating families. Geographic groups with fewer than 10 subjects were excluded from classification to ensure stable class estimates.

#### Visualisation

All plots were generated using matplotlib (v3.7+) and seaborn (v0.12+) with a consistent publication style. Figures include alpha diversity boxplots, PCoA ordination plots, differential abundance heatmaps (Z-scored per geography), volcano plots for pairwise comparisons, IGHI distribution boxplots, IGHI component heatmaps, and geography classifier confusion matrices and feature importance bar plots.

The scripts used for all statistical analysis and figure generation are available at: https://github.com/TusharParihar128/BioMeBar_Indian_Healthy_Cohorts_datasets.

## 3. Results

### 3.1 Unified matrix and cohort summary

The harmonised family-level matrix comprises 1,672 samples from 1,456 unique individuals across 380 bacterial families after NCBI family-level taxonomic harmonisation. Eight cohorts contributed subjects from 20 geographic locations spanning six Indian geographic zones (North, South, East, West, Central, Northeast) plus 61 India-based subjects from the D’Aloisio cohort without city-level metadata (designated “Indian_Unspecified”). The largest contributor was the LogMPIE cohort (n=1,004; 14 cities), followed by Jeyaram (n=214; Manipur, three seasons), Tandon (n=160; Ahmedabad, two timepoints), Dhakan (n=110; Bhopal and Kasaragod), Chaudhari (n=50; Pune), Kumbhare (n=42; Pune), Kamaldeep (n=31; Ladakh, Jaisalmer, and Khargone), and D’Aloisio (n=61; Indian residents only, diaspora excluded). Merging the Chaudhari and Kumbhare cohorts yielded a combined Pune class of 92 subjects. By zone, the West was best represented (n=443), followed by South (n=307), Northeast (n=297), North (n=200), Central (n=197), East (n=167), and Indian_Unspecified (n=61).

**Fig. 1.**
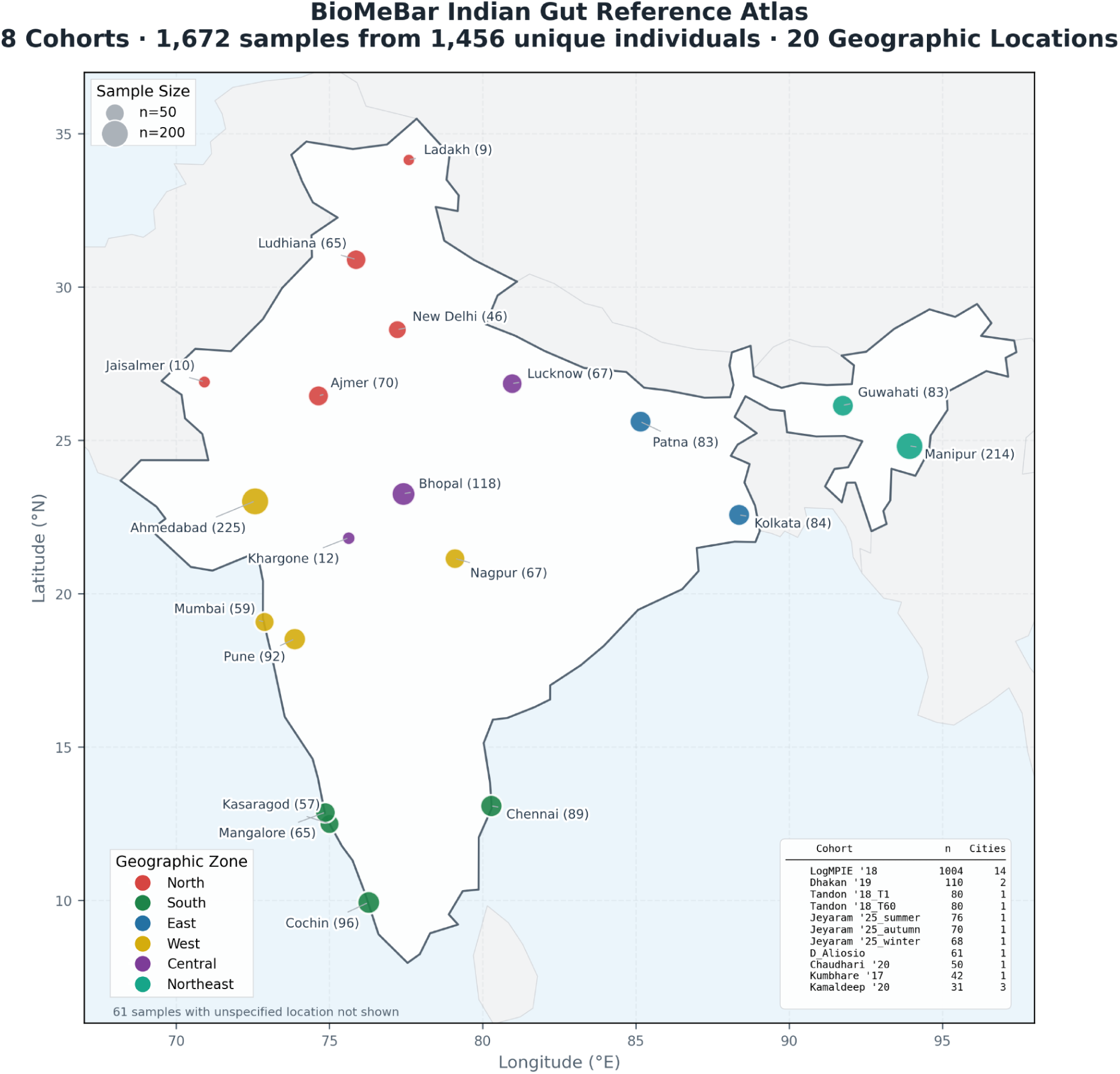
Geographic distribution of cohorts in the BioMeBar Indian Gut Reference Atlas. Circle size is proportional to √n (sample size); colour indicates geographic zone assignment. The atlas spans 20 geographic locations across six Indian zones from eight independent cohorts (n = 1,672 samples from 1,456 unique individuals). Inset table summarises cohort sample sizes. Country boundary from Natural Earth 110 m.

Seventeen families were identified as core members of the healthy Indian gut at a prevalence threshold of 80%: Lachnospiraceae (100.0%), Veillonellaceae (99.8%), Prevotellaceae (99.5%), Bacteroidaceae (98.9%), Bifidobacteriaceae (98.6%), Enterobacteriaceae (97.3%), Lactobacillaceae (96.8%), Coriobacteriaceae (96.8%), Sutterellaceae (95.7%), Streptococcaceae (93.1%), Oscillospiraceae (92.7%), Rikenellaceae (91.4%), Peptostreptococcaceae (88.0%), Erysipelotrichaceae (87.5%), Tannerellaceae (86.0%), Selenomonadaceae (84.4%), and Desulfovibrionaceae (82.8%).

**Fig. 2.**
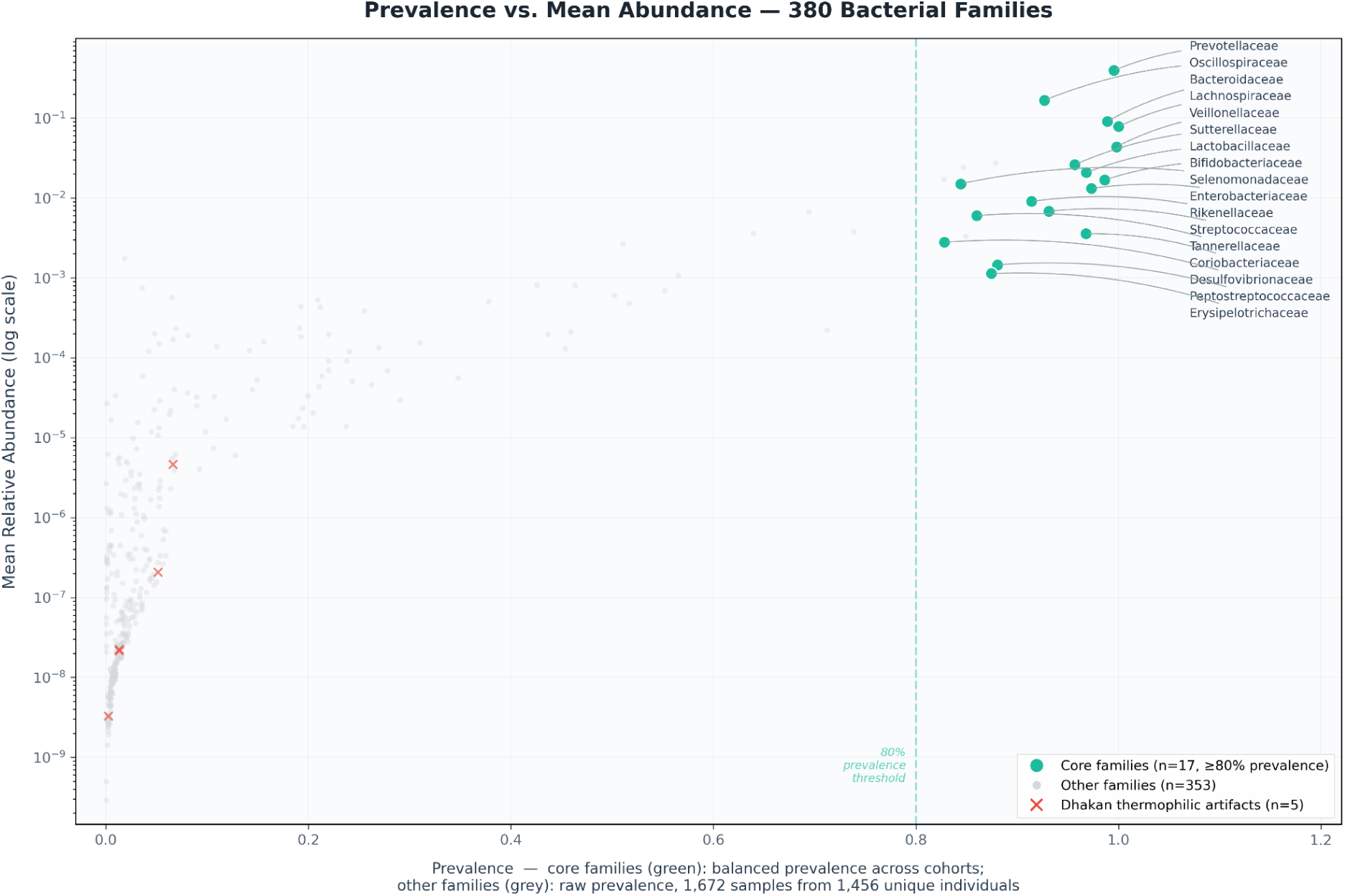
Prevalence vs. mean relative abundance for 375 bacterial families detected across the unified atlas. Each point represents one family (n = 380 total; 5 families with zero abundance in every sample are not shown, as log-scaled abundance cannot be plotted at zero). The dashed vertical line marks the 80% prevalence threshold used to define core families. For the 17 core families (green; labelled), the x-position is each family’s balanced prevalence, averaged across the eight constituent cohorts to correct for unequal cohort sizes; all other families (grey, n = 353) are plotted by raw prevalence across the full pooled dataset (1,672 samples from 1,456 unique individuals). Five families previously flagged as candidate thermophilic sequencing artifacts from the Dhakan cohort (red ×) are shown for reference. Families in the upper-right are both prevalent and abundant; those in the lower-left are rare and low-abundance.

Family accumulation analysis indicated near-saturation at approximately 375 families, suggesting the atlas captures the substantial majority of detectable family-level diversity in the Indian healthy gut at the sequencing depths represented.

### 3.2 Alpha and beta diversity across geographies

Alpha diversity differed significantly across geographic zones for all four metrics (Kruskal-Wallis, all p < 10⁻²¹). Mean Shannon diversity ranged from 1.52 ± 0.45 (West) to 1.83 ± 0.23 (Northeast), while observed family richness ranged from 26.5 ± 6.6 (North) to 63.1 ± 58.3 (Central). The high richness variance in the Central zone reflects the contribution of the Dhakan WGS cohort, which detects more families than 16S-only methods.

Beta diversity analysis by PERMANOVA on Bray–Curtis distances revealed significant compositional differences across all grouping variables (999 permutations, all p = 0.001): geography (pseudo-F = 43.4), cohort (pseudo-F = 89.6), geographic zone (pseudo-F = 52.3), and sequencing platform (pseudo-F = 146.5). PCoA ordination (PC1 = 39.0%, PC2 = 13.2%) showed partial separation by geographic zone, with Northeast samples forming the most distinct cluster; however, restricted-permutation PERMANOVA showed that zone does not explain variance beyond cohort (restricted p = 0.896), indicating that the Northeast cluster reflects the single Jeyaram cohort rather than an independent geographic signal. City-level geography, by contrast, did explain variance beyond cohort (restricted p = 0.001).

**Fig. 3.**
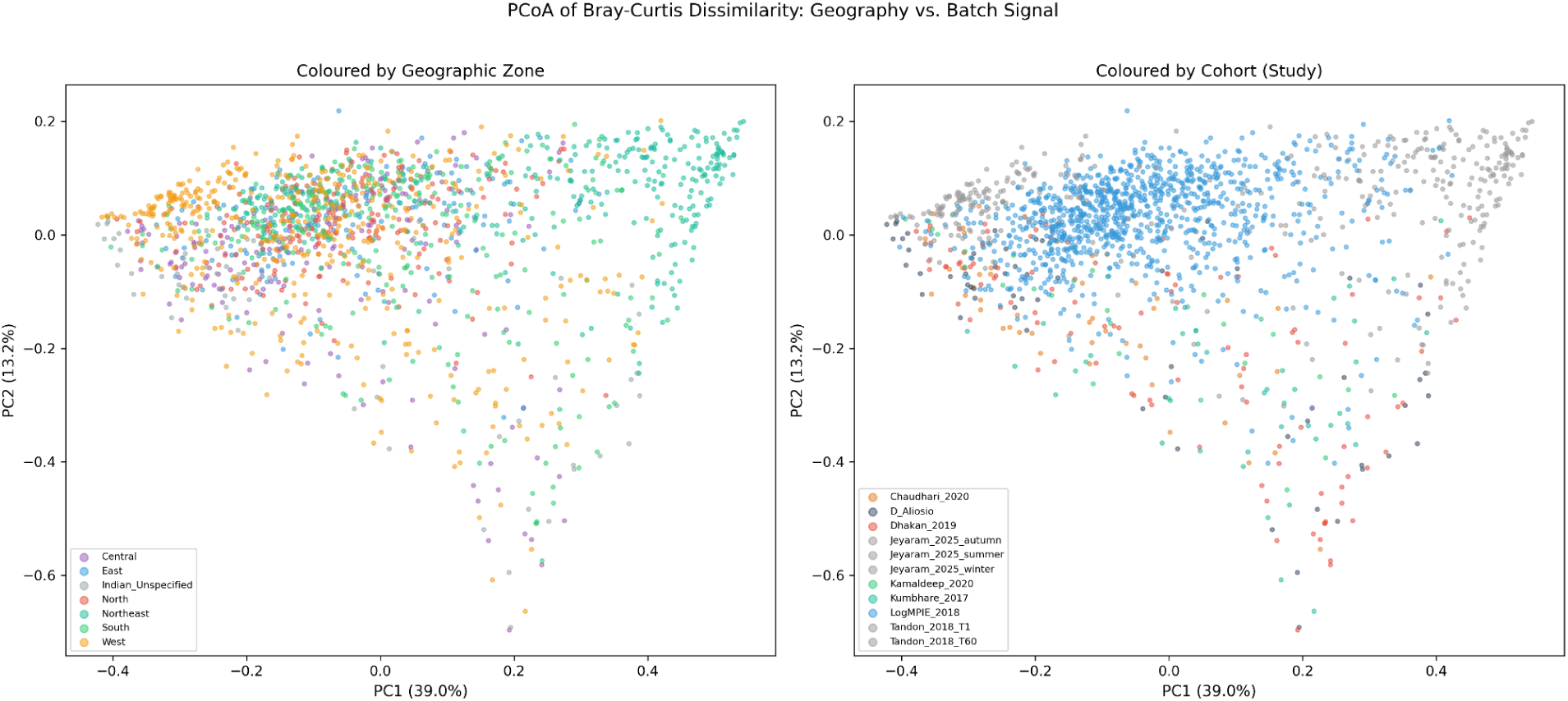
Principal Coordinates Analysis (PCoA) of Bray–Curtis dissimilarities among 1,672 samples from 1,456 unique individuals, coloured by (a) geographic zone (7 zones) and (b) cohort identity (11 cohort labels, including separate timepoints for the Tandon and Jeyaram longitudinal studies). PC1 explains 39.0% and PC2 explains 13.2% of total variance. Samples from the Northeast zone - drawn entirely from a single cohort (Jeyaram 2025)[15] - form the most visually distinct cluster in both panels; restricted-permutation PERMANOVA indicates this reflects cohort/batch structure rather than an independent geographic effect (zone beyond cohort: restricted p = 0.896), whereas city-level geography does explain variance beyond cohort (restricted p = 0.001). The visually comparable separation by cohort and zone in (a) and (b) illustrates the confounded study–geography structure of the atlas.

Critically, cohort identity explained comparable variance to geography (R² = 35.0% vs. 34.5%), reflecting the confounded study design in which 20 of 22 city-level geographies were sampled by a single study. Variance partitioning decomposed total explained variance into pure geography (4.1%), pure cohort (4.7%), and shared/confounded (30.4%), with 60.9% residual. PERMDISP confirmed significant heterogeneity of multivariate dispersion across geographies (Kruskal-Wallis H = 515.6, p < 10^-96) and cohorts (H = 344.3, p < 10^-68), indicating that PERMANOVA results reflect both location and dispersion differences.

**Fig. 4.**
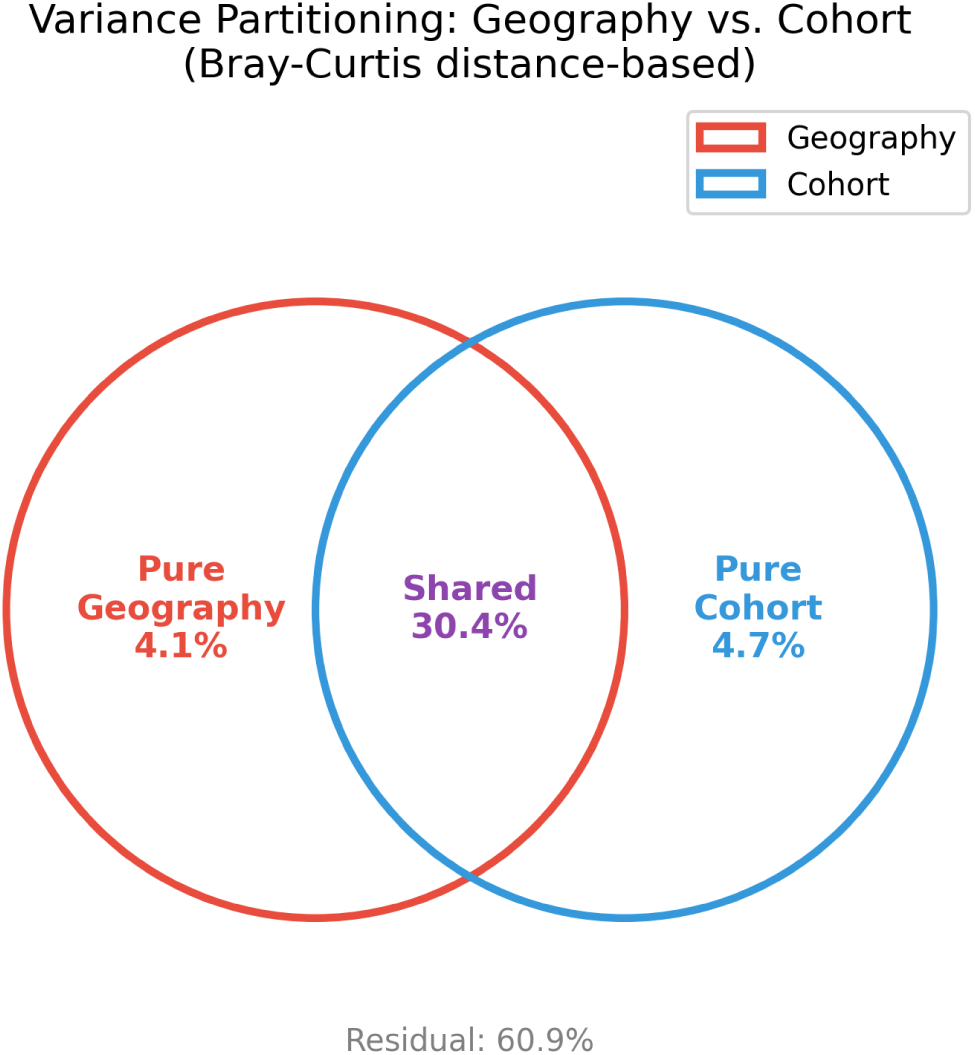
Variance partitioning of Bray–Curtis dissimilarity into pure geography (4.1%), pure cohort (4.7%), shared/confounded (30.4%), and residual (60.9%) components. The large shared fraction indicates that geographic and cohort effects cannot be cleanly separated in this multi-study design.

Restricted permutation PERMANOVA, in which geography labels were shuffled only within cohorts, confirmed that city-level geography remains significant after controlling for batch (p = 0.001). However, zone-level geography did not survive this correction (p = 0.896), indicating that much of the between-zone compositional variance is attributable to between-study differences. Mantel test results were consistent: geographic distance correlated weakly but significantly with compositional distance (n = 1,611 samples from 1,395 unique subjects (after excluding 61 D’Aloisio Indian_Unspecified) with city-level coordinates; Spearman r = 0.17, p = 0.001), but this correlation disappeared within the LogMPIE cohort alone (r = 0.009, p = 0.44), where 14 cities were sampled using identical protocols.

### 3.3 Community types

Clustering analysis on PCoA-projected Bray-Curtis distances identified two optimal community types (k-medoids, silhouette = 0.343): a Prevotellaceae-dominant type (n = 1,153, 69%) characterised by high Prevotellaceae (mean relative abundance 0.515), Oscillospiraceae (0.132), and Bacteroidaceae (0.079); and an Oscillospiraceae-dominant type (n = 519, 31%) characterised by Oscillospiraceae (0.244), Lachnospiraceae (0.149), and Prevotellaceae (0.126). The Oscillospiraceae-dominant type was enriched in Northeast zone subjects (40% of cluster), while the Prevotellaceae-dominant type was distributed more evenly across zones, with West subjects comprising 31%.

**Fig. 5.**
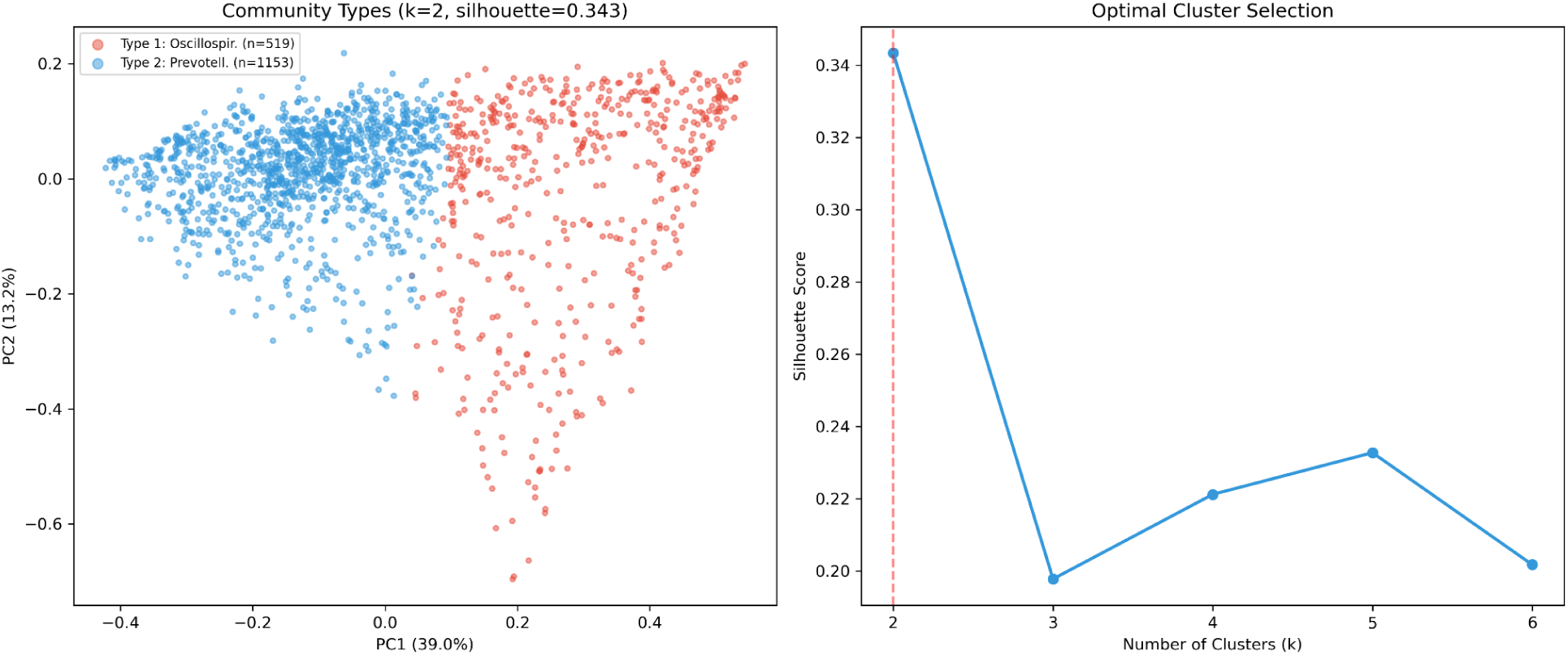
Two Indian gut community types identified by k-means clustering (k = 2, silhouette = 0.343) on Bray–Curtis distances. (a) PCoA ordination (PC1 39.0%, PC2 13.2%) coloured by community-type assignment: a Prevotellaceae-dominant type (n = 1,153, 69%) and an Oscillospiraceae-dominant type (n = 519, 31%). (b) Silhouette score across candidate cluster numbers (k = 2–6), confirming k = 2 as optimal.

This two-type structure contrasts with the three-enterotype model (Prevotella, Bacteroides, Ruminococcus) commonly reported in Western populations, consistent with the higher prevalence of Prevotella-dominant profiles in Indian populations linked to plant-rich, high-fibre diets.

### 3.4 IGHI distribution and geography-stratified norms

The Indian Gut Health Index (IGHI) across all 1,672 samples from 1,456 unique individuals showed a mean of 71.75 ± 10.57, with geography-stratified zone means ranging from 67.6 ± 10.9 (Indian_Unspecified) to 72.9 ± 11.3 (Central). The inter-zone range was narrow (5.3 points), indicating that the geography-stratified percentiling within the IGHI design effectively normalises regional baseline differences.

**Fig. 6.**
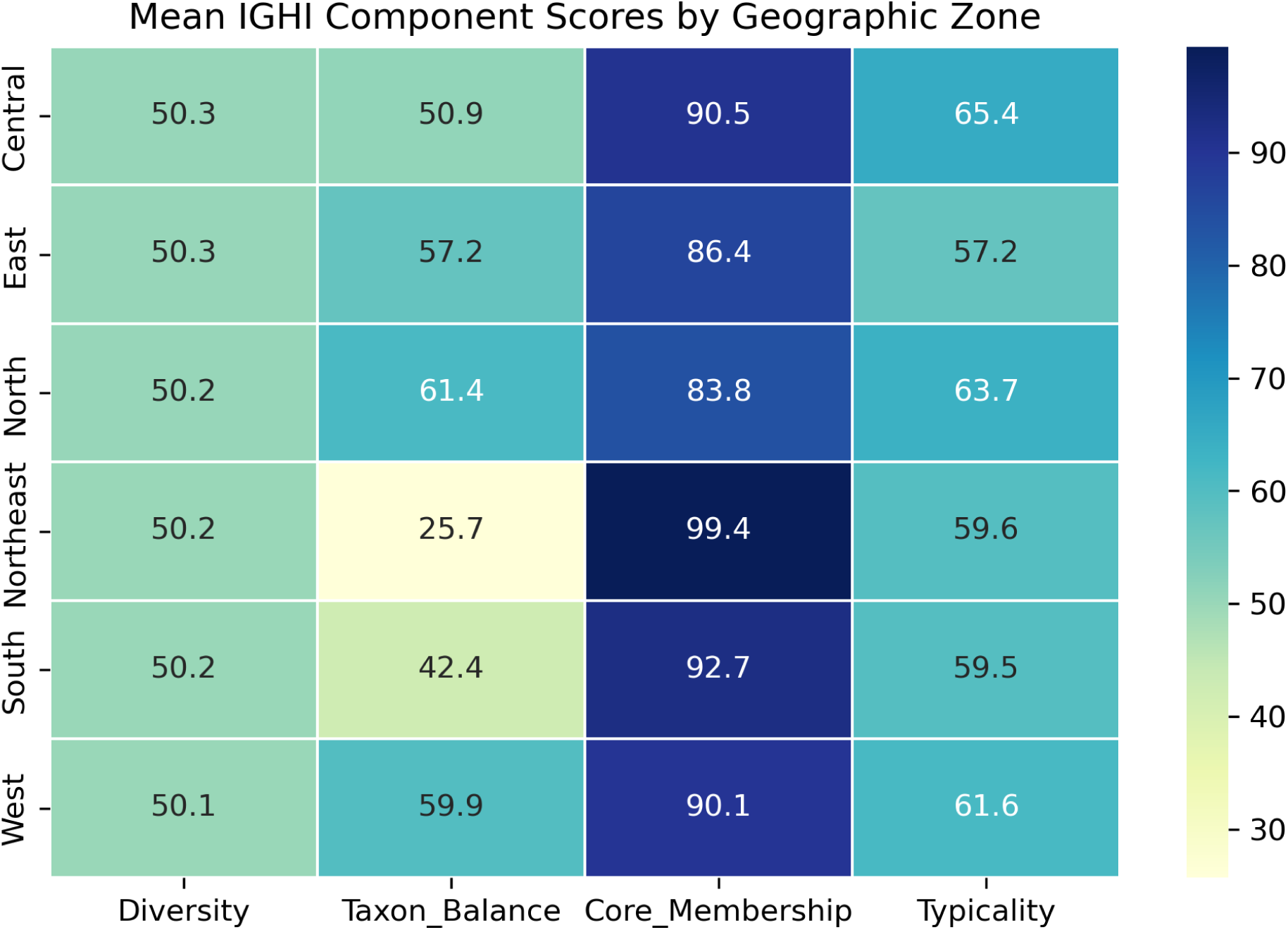
IGHI component scores by geographic zone. Each row is a zone; columns show the four component scores (Diversity, Taxon Balance, Core Membership, Community Typicality). Values are zone means. The narrow inter-zone IGHI range (5.3 points) reflects the geography-stratified percentiling design.

IGHI component analysis revealed that the four components capture largely independent dimensions of gut health. Spearman correlations between components were low for most pairs (|ρ| < 0.3), with the exception of Diversity vs. Taxon Balance (ρ = -0.41) and Taxon Balance vs. Core Membership (ρ = **−0.32**). The negative Diversity-Balance correlation indicates that more diverse gut communities do not necessarily have higher beneficial-to-opportunistic family ratios, suggesting these components capture complementary information.

Weight sensitivity analysis across six alternative weighting schemes (equal weights, diversity-heavy, balance-heavy, typicality-heavy, core-heavy) showed high rank stability: the default (30/25/20/25) scheme correlated at Spearman ρ ≥ 0.88 with all alternatives except the balance-heavy scheme (ρ = 0.72), confirming that IGHI rankings are robust to moderate weight perturbation.

**Fig. 7.**
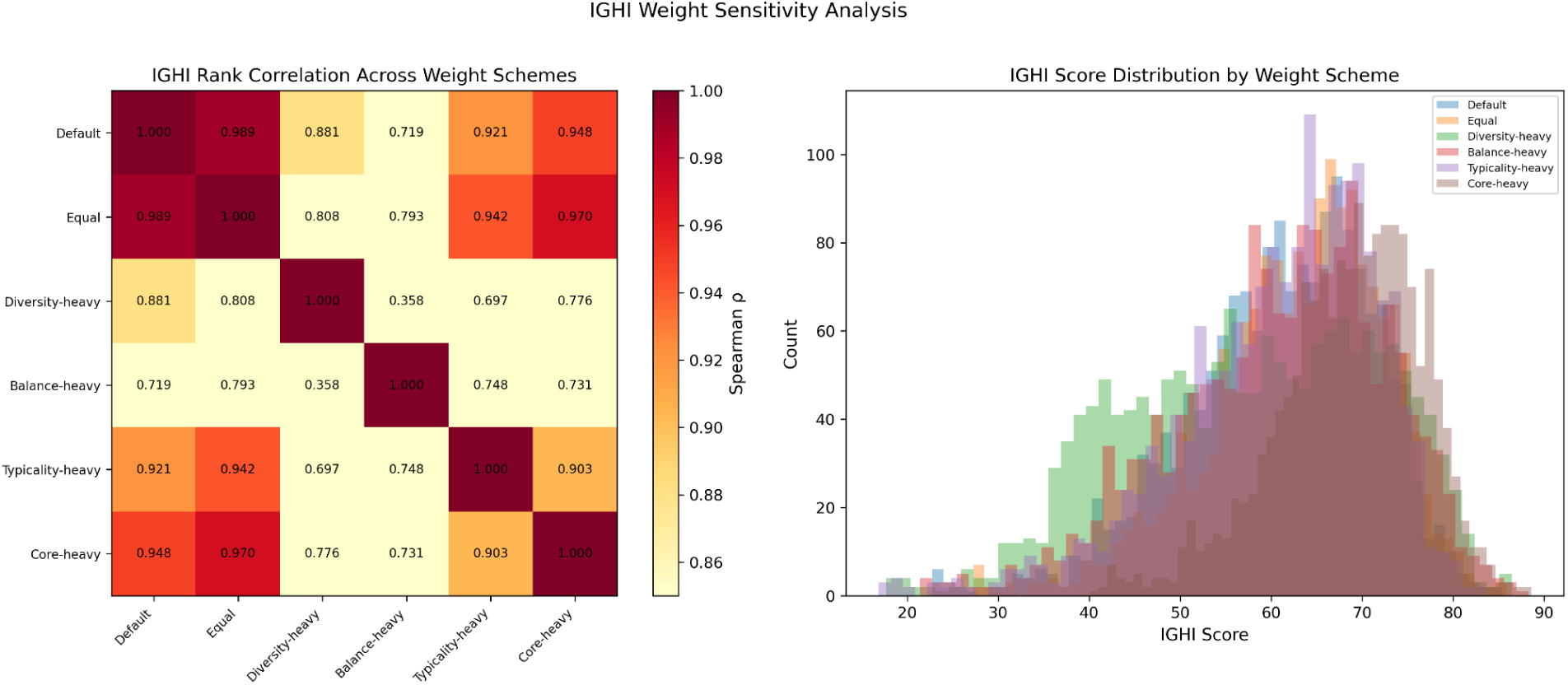
IGHI weight sensitivity analysis. (a) Pairwise Spearman rank correlations (ρ) of composite IGHI scores under six weighting schemes: Default (Diversity 0.30, Taxon Balance 0.25, Core Membership 0.20, Typicality 0.25), Equal (0.25 each), and four schemes each up-weighting one component to 0.50, with the remaining weight split evenly across the other three (Diversity-heavy, Balance-heavy, Typicality-heavy, Core-heavy). Relative to the default scheme, four of the five alternative schemes show high agreement (ρ = 0.881–0.989); Balance-heavy shows weaker agreement (ρ = 0.719). Across all pairwise comparisons, correlations range as low as ρ = 0.358 (Diversity-heavy vs. Balance-heavy), indicating that rankings are sensitive to simultaneous, opposing extreme reweighting of these two components. (b) Distribution of IGHI scores (n = 1,672 samples from 1,456 unique individuals) under each weighting scheme.

### 3.5 Differentially abundant families

Kruskal-Wallis testing identified the majority of families as significantly differentially abundant across geographic zones (FDR q < 0.05). Indicator species analysis (IndVal)[28] identified 319 of 380 families as significant zone indicators, with the Northeast zone contributing the strongest indicators: *Streptococcaceae* (IndVal = 0.748), *Budviciaceae* (0.721), *Oceanospirillaceae* (0.714), *Peptococcaceae* (0.712), and *Anaerovoracaceae* (0.711). The West zone was indicated by *Butyricicoccaceae* (0.520) and *Succinivibrionaceae* (0.498); the South by *Pasteurellaceae* (0.306) and *Sutterellaceae* (0.250); the East by *Treponemataceae* (0.435) and *Desulfovibrionaceae* (0.370); and the Central zone by *Enterobacteriaceae* (0.332) and *Bacteroidaceae* (0.228).

**Fig. 8.**
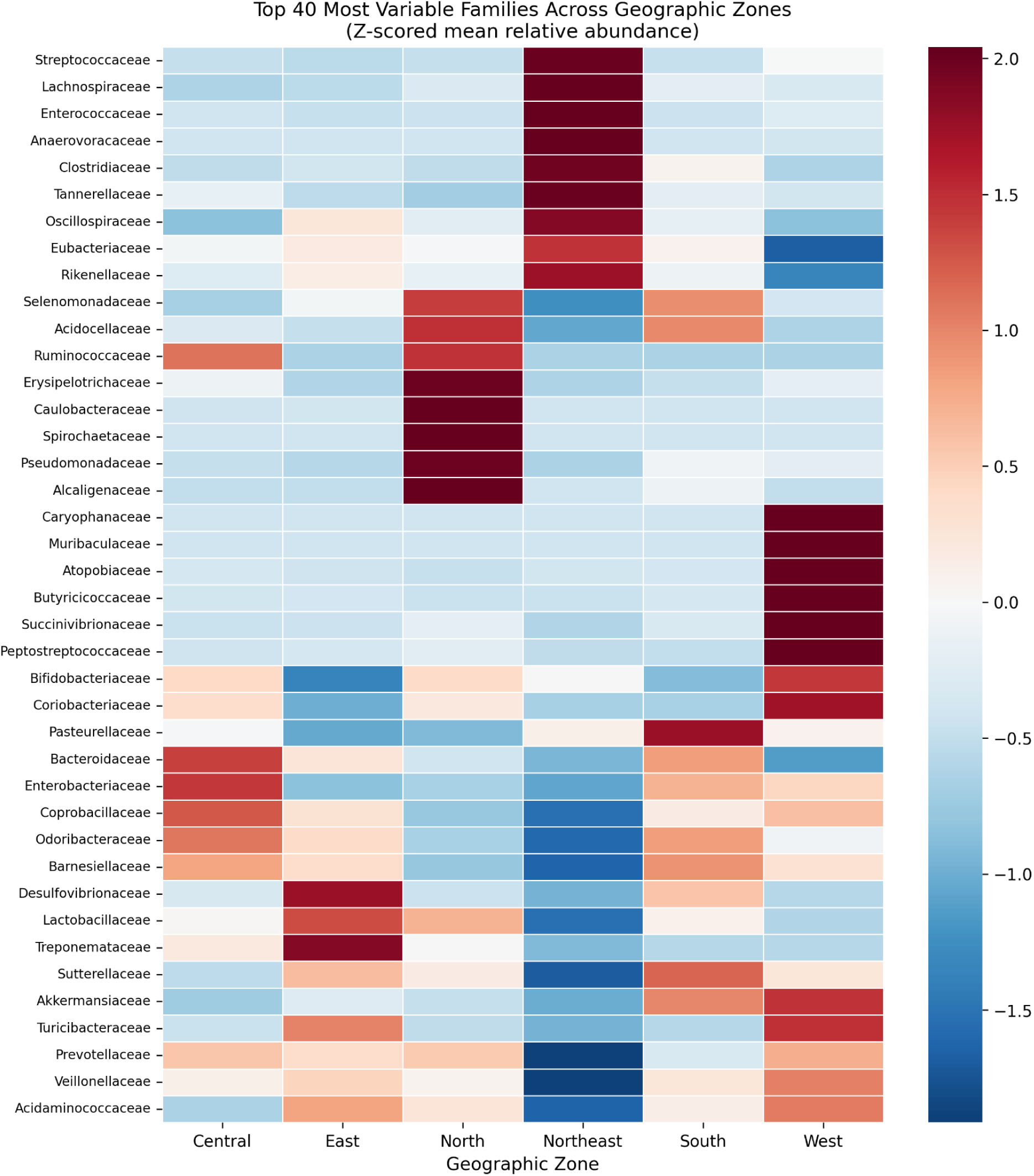
Differential abundance heatmap of the top 40 most variable families across geographic zones. Colour intensity represents Z-scored mean relative abundance within each zone (red = enriched, blue = depleted relative to the all-zone mean). Families are clustered by similarity of their zone abundance profiles. Asterisks indicate significance (Kruskal–Wallis FDR q < 0.05).

**Fig. 9.**
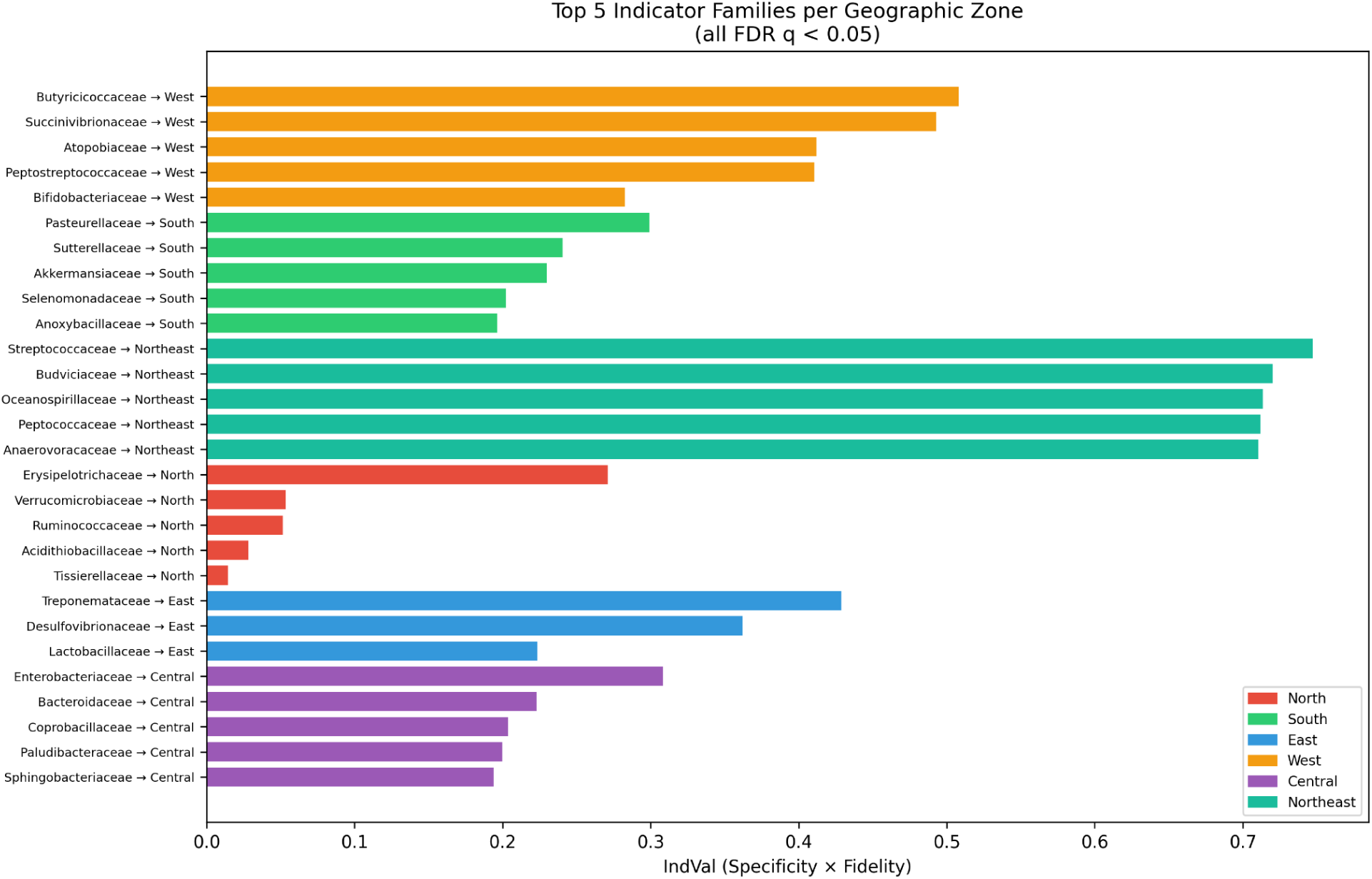
Top indicator families per geographic zone by IndVal score. Bar length represents the IndVal statistic (0–1); colour indicates the zone for which each family is an indicator. The Northeast zone contributes the strongest indicators (Streptococcaceae, IndVal = 0.748), consistent with its distinct dietary and geographic profile.

The *Prevotella/Bacteroides* ratio showed a strong zone gradient, with Northeast samples being overwhelmingly Prevotella-dominant and West samples showing the most balanced ratio, consistent with the dietary gradient from the fermented-food-rich diets of Manipur to the diverse urban diets of western Indian cities.

Of the 300 pairwise zone comparisons for the top 20 most variable families, 216 (72%) were significant after FDR correction, with 56 (19%) showing large effect sizes (Cliff’s delta |δ| > 0.474).

### 3.6 Geography classifier performance

A Random Forest classifier trained on family-level abundance profiles to predict geographic zone achieved 68.4% balanced accuracy under subject-grouped stratified 5-fold cross-validation, substantially exceeding the permutation null baseline of 16.6% ± 1.2% (n = 20 permutations). Feature selection with the top 75 families achieved the highest balanced accuracy (69.1%). The most discriminating families by Gini importance were Treponemataceae (0.038), Eubacteriaceae (0.037), Pasteurellaceae (0.031), and Streptococcaceae (0.026).

Per-zone performance varied: Northeast achieved the highest F1 score (0.92, precision 0.99, recall 0.86), followed by West (F1 = 0.80, precision 0.89, recall 0.73). Central zone showed high precision (0.86) but low recall (0.34), suggesting its compositional profile overlaps with other zones.

Leave-One-Study-Out (LOSO) cross-validation produced mixed results: balanced accuracy was 1.0 when holding out Chaudhari, Kumbhare, or Tandon (all from the West zone, where LogMPIE provides training data from the same geography), but dropped to 0.0 when holding out Dhakan, Jeyaram, or Kamaldeep (which contribute unique geographies not represented in other studies). This pattern confirms that the classifier’s performance is partly attributable to within-geography cross-study generalisation and partly to the confounded study-geography structure of the atlas.

## 4. Discussion

### 4.1 The need for an Indian-specific reference

This work addresses a fundamental gap in gut microbiome science: the absence of a harmonised healthy reference for the Indian population. Existing reference databases - HMP (n=242, predominantly US), MetaHIT (n=124, European), the Flemish Gut Flora Project (n=1,106, Belgian), and the American Gut (n>10,000, primarily US/UK) were constructed from populations with substantially different diets, genetics, and environmental exposures than those prevailing across India. The Gut Microbiome Health Index [29] represents an important step toward a universal health-associated scoring model, but its training data over-represents Western cohorts and its taxon-level features may not weight Indian-prevalent taxa appropriately. Our atlas provides the Indian-specific calibration data that such models require.

The geographic variation we document within India underscores why a single national reference is insufficient. India’s dietary landscape - from the strictly vegetarian communities of Gujarat and Rajasthan to the fermented-food-rich diets of the Northeast, the seafood diets of Kerala, and the wheat-and-dairy staples of the North creates microbiome gradients as large as those observed between countries in international studies. The IGHI’s geography-stratified normalization addresses this by comparing each individual to their regional healthy baseline rather than to a pan-Indian average that may not represent any actual community.

### 4.2 Harmonisation at family level: rationale and limitations

Our decision to harmonise at family rank represents a deliberate trade-off. Genus- or species-level resolution would preserve finer ecological detail, but the heterogeneity of input classifiers makes sub-family harmonisation unreliable across cohorts. GreenGenes 13.8 and SILVA 138.1 differ in genus assignments for 15–20% of sequences [19], [30]; MetaPhlAn and OTU-based classifiers operate on fundamentally different principles (clade-specific markers vs. sequence similarity clustering); and HITChip probe-based groups do not correspond exactly to conventional taxonomic ranks. Family-level aggregation absorbs these discrepancies while preserving the distinctions most relevant to gut health assessment: *Lachnospiraceae* vs. *Enterobacteriaceae*, *Bifidobacteriaceae* vs. *Clostridiaceae* - and to the functional inferences that inform probiotic and dietary interventions.

The NCBI Taxonomy backbone was chosen over GreenGenes (unmaintained since 2013) and the SILVA taxonomy (internally consistent but less widely adopted in clinical and commercial contexts). The resolution cascade: exact match, synonym lookup, genus-to-family fallback, and explicit “Unassigned” flagging ensures transparent traceability: every taxonomic mapping decision is recorded and deposited with the dataset.

A known limitation of family-level analysis is the loss of within-family diversity. Lachnospiraceae, for example, encompasses both butyrate-producing genera (*Roseburia*, *Coprococcus*) and genera with less clear functional roles. Genus-level analysis within individual cohorts can supplement the family-level atlas where per-cohort data quality permits.

### 4.3 The IGHI as a clinical and commercial scoring model

The IGHI was designed to fill a practical gap: no existing gut health scoring model is calibrated for Indian populations. Commercial gut tests sold in India currently report against Western reference ranges, and clinical interpretation relies on literature derived from non-Indian cohorts. The IGHI’s four-component structure captures complementary dimensions of gut health:

- **Diversity** reflects ecosystem stability and resilience, the most replicated single correlate of gut health across populations [31].
- **Taxon balance** captures the ratio of families consistently associated with health vs. dysbiosis, providing a functional summary complementary to raw diversity.
- **Core membership** ensures that the fundamental Indian-specific community structure is intact - an individual missing core families is, by definition, deviating from the healthy Indian norm regardless of their diversity or balance scores.
- **Community typicality** positions the individual within the multidimensional space of healthy profiles from their region, capturing compositional features not reducible to univariate metrics.

The default weight allocation (diversity 0.30, taxon balance 0.25, core membership 0.20, typicality 0.25) is a starting point informed by the relative robustness and interpretability of each component. Formal optimisation of weights, for example, using supervised learning against labelled healthy/disease cohorts - is a priority for future work, particularly as Indian IBS and IBD sequencing datasets become available.

We emphasise that the preliminary interpretive thresholds (Excellent: 80–100, Good: 60–79, Moderate: 40–59, Below average: 20–39, Low: 0–19) are derived from the healthy atlas distribution and have not been validated against disease outcomes. The IGHI should be considered a research-grade scoring framework requiring prospective clinical validation before deployment in diagnostic or clinical-decision contexts.

### 4.4 Comparison with prior Indian and global microbiome resources

The closest precedent to this work is the [8] study, which remains the largest single-cohort Indian gut microbiome characterisation with integrated metabolomics. Our atlas extends the Dhakan cohort in three directions: geographic coverage (20 regions vs. 2), sample size (approximately 1,672 samples from 1,456 unique individuals vs. 110 subjects), and the provision of a unified analytical framework (the IGHI) for scoring individual profiles against the reference.

The HRGM (Kim et al., 2021)[6] provides a genome-level catalogue of 232,098 non-redundant genomes from 51,316 samples, but its primary value is as a reference for metagenome-assembled genome analysis rather than a scoring model for individual health assessment. The GMHI [29] is architecturally closest to the IGHI: both are composite scores derived from taxonomic abundance profiles but GMHI was trained predominantly on Western cohorts and operates at species level, limiting its applicability to Indian populations characterised at lower taxonomic resolution.

Bhute et al. (2016)[12], one of the earliest Indian gut microbiome studies, characterised approximately 34 subjects from Pune but deposited limited raw data (8 samples in NCBI SRA, with additional data in the DDBJ DRA archive), precluding inclusion in our atlas at scale.

### 4.5 Limitations and future directions

Several limitations warrant acknowledgement. First, the atlas is constructed exclusively from healthy subjects; its clinical utility depends on validation against Indian disease cohorts.

Second, the use of family-level taxonomy sacrifices ecological resolution. Future iterations of the atlas, incorporating whole-genome shotgun data from the D’Aloisio cohort (110 WGS samples, processed through KneadData v3.1, MetaPhlAn 4.2.5, and HUMAnN 3.9) and any newly published Indian cohorts, will support genus, or species-level analysis where input data quality permits. The D’Aloisio cohort uniquely contributes both 16S rRNA and WGS data for the same 110 subjects, enabling future cross-platform validation of the 16S-based taxonomic profiles and preliminary functional profiling via pathway and gene family abundance.

Repeated measures from 152 individuals introduced non-independence; a sensitivity analysis using one sample per subject confirmed that results were robust to this structure.

Third, geographic coverage remains uneven: western India (Pune, Ahmedabad) is over-represented relative to the populous northern plains (Uttar Pradesh, Bihar) and southern India (Tamil Nadu, Karnataka). Expanding coverage to these underrepresented regions is a priority.

Fourth, the IGHI weights are heuristic rather than optimised. Once labelled healthy/disease training data become available, supervised weight optimisation (e.g., logistic regression of IGHI components against health status) should replace the current expert-assigned weights.

Fifth, batch effects from different sequencing platforms and bioinformatics pipelines cannot be fully eliminated by family-level harmonisation. The leave-one-cohort-out cross-validation in the geography classifier provides an empirical test of the extent to which batch effects confound biological signals.

Finally, the metabolomics layers available from the Dhakan (GC-MS fecal + serum) and Jeyaram (LC-HRMS fecal) cohorts are not directly comparable due to different analytical platforms and metabolite identification approaches. Integration of microbiome and metabolome data will require platform-specific normalization and should be treated as exploratory.

Despite these limitations, the BioMeBar^TM^ Indian Gut Reference Atlas represents the most comprehensive harmonised healthy gut microbiome resource currently available for the Indian population, and the IGHI provides the first population-calibrated framework for Indian gut health scoring. We anticipate that this resource will not only help calibrate new reference ranges for Indian gut tests but also support the development of India-specific probiotics, prebiotics, postbiotics, synbiotics, and personalized nutritional interventions grounded in the actual composition of the healthy Indian gut.

## 5. Data Availability

The harmonised family-level abundance matrix, NCBI taxonomy mapping tables, per-cohort processing documentation, IGHI scoring implementation, and all analysis scripts are available at https://github.com/TusharParihar128/BioMeBar_Indian_Healthy_Cohorts_datasets. Raw sequence data for individual cohorts are deposited in NCBI SRA under the BioProject accessions listed in Table 1. The Jeyaram cohort data are available from Zenodo (DOI: 10.5281/zenodo.14424024). The Dhakan cohort supplementary data are available from GigaDB (DOI: 10.5524/100548). The IGHI scoring code and Phase 2 analysis pipeline are available at https://github.com/TusharParihar128/BioMeBar_Indian_Healthy_Cohorts_datasets.

## 6. Author Contributions

N.D. conceived and directed the study, defined the IGHI scoring model, and wrote the manuscript. T.P. performed literature identification, dataset acquisition, SRA processing, taxonomic harmonisation, and implemented the analysis pipeline. Both authors reviewed and approved the final manuscript. R.M. helped finalize python scripts.

## 9. Acknowledgements

We thank the original investigators of the eight included cohorts for making their data publicly available, without which this harmonisation would not have been possible. We acknowledge NCBI, SILVA, DADA2 and the QIIME 2 development team for the bioinformatics infrastructure underpinning this work.

## References

[1] C. Huttenhower et al., “Structure, function and diversity of the healthy human microbiome,” Nature, vol. 486, no. 7402, pp. 207–214, Jun. 2012, doi: 10.1038/nature11234.

[2] J. Qin et al., “A human gut microbial gene catalogue established by metagenomic sequencing,” Nature, vol. 464, no. 7285, pp. 59–65, 2010, doi: 10.1038/nature08821.

[3] G. Falony et al., “Population-level analysis of gut microbiome variation,” Science (1979)., vol. 352, no. 6285, pp. 560–564, Apr. 2016, doi: 10.1126/science.aad3503.

[4] M. Daniel et al., “American Gut: an Open Platform for Citizen Science Microbiome Research,” mSystems, vol. 3, no. 3, pp. 10.1128/msystems.00031-18, May 2018, doi: 10.1128/msystems.00031-18.

[5] A. Almeida et al., “A unified catalog of 204,938 reference genomes from the human gut microbiome,” Nat. Biotechnol., vol. 39, no. 1, pp. 105–114, Jan. 2021, doi: 10.1038/s41587-020-0603-3.

[6] C. Y. Kim et al., “Human reference gut microbiome catalog including newly assembled genomes from under-represented Asian metagenomes,” Genome Med., vol. 13, no. 1, Dec. 2021, doi: 10.1186/s13073-021-00950-7.

[7] R. J. Abdill, E. M. Adamowicz, and R. Blekhman, “Public human microbiome data are dominated by highly developed countries,” PLoS Biol., vol. 20, no. 2, Feb. 2022, doi: 10.1371/journal.pbio.3001536.

[8] D. B. Dhakan et al., “The unique composition of Indian gut microbiome, gene catalogue, and associated fecal metabolome deciphered using multi-omics approaches,” Gigascience, vol. 8, no. 3, Jan. 2019, doi: 10.1093/gigascience/giz004.

[9] K. Kaur, I. Khatri, A. Akhtar, S. Subramanian, and T. N. C. Ramya, “Metagenomics analysis reveals features unique to Indian distal gut microbiota,” PLoS One, vol. 15, no. 4, Apr. 2020, doi: 10.1371/journal.pone.0231197.

[10] A. K. Dubey et al., “LogMPIE, pan-India profiling of the human gut microbiome using 16S rRNA sequencing,” Sci. Data, vol. 5, 2018, doi: 10.1038/sdata.2018.232.

[11] D. Tandon et al., “A snapshot of gut microbiota of an adult urban population from Western region of India,” PLoS One, vol. 13, no. 4, Apr. 2018, doi: 10.1371/journal.pone.0195643.

[12] S. Bhute et al., “Molecular characterization and meta-analysis of gut microbial communities illustrate enrichment of prevotella and megasphaera in Indian subjects,” Front. Microbiol., vol. 7, no. MAY, 2016, doi: 10.3389/fmicb.2016.00660.

[13] S. V. Kumbhare et al., “A cross-sectional comparative study of gut bacterial community of Indian and Finnish children,” Sci. Rep., vol. 7, no. 1, Dec. 2017, doi: 10.1038/s41598-017-11215-y.

[14] D. S. Chaudhari et al., “Gut, oral and skin microbiome of Indian patrilineal families reveal perceptible association with age,” Sci. Rep., vol. 10, no. 1, Dec. 2020, doi: 10.1038/s41598-020-62195-5.

[15] K. Jeyaram et al., “Fermented foods affect the seasonal stability of gut bacteria in an Indian rural population,” Nature Communications, vol. 16, no. 1, Dec. 2025, doi: 10.1038/s41467-025-56014-6.

[16] L. D. D’Aloisio et al., “The adoption of a westernized gut microbiome in Indian Immigrants and Indo-Canadians is associated with dietary acculturation,” NPJ Biofilms Microbiomes, vol. 11, no. 1, Dec. 2025, doi: 10.1038/s41522-025-00778-8.

[17] E. Bolyen et al., “Reproducible, interactive, scalable and extensible microbiome data science using QIIME 2,” Aug. 01, 2019, Nature Publishing Group. doi: 10.1038/s41587-019-0209-9.

[18] B. J. Callahan, P. J. McMurdie, M. J. Rosen, A. W. Han, A. J. A. Johnson, and S. P. Holmes, “DADA2: High-resolution sample inference from Illumina amplicon data,” Nat. Methods, vol. 13, no. 7, pp. 581–583, Jun. 2016, doi: 10.1038/nmeth.3869.

[19] C. Quast et al., “The SILVA ribosomal RNA gene database project: Improved data processing and web-based tools,” Nucleic Acids Res., vol. 41, no. D1, Jan. 2013, doi: 10.1093/nar/gks1219.

[20] G. M. Douglas et al., “PICRUSt2 for prediction of metagenome functions,” Jun. 01, 2020, Nature Research. doi: 10.1038/s41587-020-0548-6.

[21] P. Barbera et al., “EPA-ng: Massively Parallel Evolutionary Placement of Genetic Sequences,” Mar. 01, 2019, Oxford University Press. doi: 10.1093/sysbio/syy054.

[22] Y. Ye and T. G. Doak, “A parsimony approach to biological pathway reconstruction/inference for genomes and metagenomes,” PLoS Comput. Biol., vol. 5, no. 8, 2009, doi: 10.1371/journal.pcbi.1000465.

[23] C. E. Shannon, “A Mathematical Theory of Communication,” Bell System Technical Journal, vol. 27, no. 3, pp. 379–423, Jul. 1948, doi: 10.1002/j.1538-7305.1948.tb01338.x.

[24] J. R. Bray and J. T. Curtis, “An Ordination of the Upland Forest Communities of Southern Wisconsin,” Ecol. Monogr., vol. 27, no. 4, pp. 325–349, Oct. 1957, doi: 10.2307/1942268.

[25] M. J. Anderson, “A new method for non-parametric multivariate analysis of variance,” Austral Ecol., vol. 26, no. 1, pp. 32–46, Feb. 2001, doi: 10.1111/j.1442-9993.2001.01070.pp.x.

[26] N. Cliff, “Dominance Statistics: Ordinal Analyses to Answer Ordinal Questions,” Psychol. Bull., vol. 114, pp. 494–509, Nov. 1993, doi: 10.1037/0033-2909.114.3.494.

[27] L. Breiman, “Random Forests,” Mach. Learn., vol. 45, no. 1, pp. 5–32, Oct. 2001, doi: 10.1023/A:1010933404324.

[28] M. Dufrêne and P. Legendre, “SPECIES ASSEMBLAGES AND INDICATOR SPECIES:THE NEED FOR A FLEXIBLE ASYMMETRICAL APPROACH,” Ecol. Monogr., vol. 67, no. 3, pp. 345–366, Aug. 1997, doi: 10.1890/0012-9615(1997)067[0345:SAAIST]2.0.CO;2.

[29] V. K. Gupta et al., “A predictive index for health status using species-level gut microbiome profiling,” Nature Communications, vol. 11, no. 1, Dec. 2020, doi: 10.1038/s41467-020-18476-8.

[30] D. McDonald et al., “An improved Greengenes taxonomy with explicit ranks for ecological and evolutionary analyses of bacteria and archaea,” ISME Journal, vol. 6, no. 3, pp. 610–618, Mar. 2012, doi: 10.1038/ismej.2011.139.

[31] C. A. Lozupone, J. I. Stombaugh, J. I. Gordon, J. K. Jansson, and R. Knight, “Diversity, stability and resilience of the human gut microbiota,” Sep. 13, 2012. doi: 10.1038/nature11550.

