## Supplementary for "BioMeBar Indian Gut Reference Atlas: A Harmonised Multi-Cohort Healthy Microbiome Baseline and the Indian Gut Health Index"

### Supplementary Methods

**S.M.1. Restricted Permutation PERMANOVA**

To disentangle geographic signal from cohort/batch effects, we performed PERMANOVA with permutations restricted within cohort strata. For each of 999 permutations, geography labels were shuffled only among subjects within the same cohort, ensuring that the null distribution reflects within-cohort geographic variation only. This was performed at both city level (22 geographies) and zone level (7 zones including Indian_Unspecified). Implementation: custom permutation scheme using scikit-bio's permanova function with manually constructed permutation indices.

**S.M.2. PERMDISP (Multivariate Dispersion Test)**

For each grouping variable (geography, cohort, zone, platform), we computed each subject's distance to their group centroid in Bray–Curtis space, then tested for differences in dispersion across groups using the Kruskal–Wallis H test. Significant PERMDISP indicates that PERMANOVA results may partly reflect dispersion (heteroscedasticity) rather than location differences. Implementation: per-group centroid distances computed from the full Bray–Curtis distance matrix.

**S.M.3. Variance Partitioning**

Total explained variance in the Bray–Curtis distance matrix was decomposed into pure geography, pure cohort, shared (confounded), and residual components:

$R^2$_geo: PERMANOVA $R^2$ for geography alone
$R^2$_cohort: PERMANOVA $R^2$ for cohort alone
$R^2$_both: PERMANOVA $R^2$ for geography + cohort (additive model)

Pure geography = $R^2$_both − $R^2$_cohort
Pure cohort = $R^2$_both − $R^2$_geo
Shared = $R^2$_geo + $R^2$_cohort − $R^2$_both
Residual = 1 − $R^2$_both

**S.M.4. Indicator Species Analysis (IndVal)**

The IndVal statistic [28] was computed for each family × zone combination as IndVal = specificity × fidelity, where specificity is the proportion of total abundance of that family found in the target zone, and fidelity is the proportion of samples in the target zone where the family is present. Significance was assessed by 999 permutations.

**S.M.5. Community Type (Enterotype) Clustering**

k-medoids clustering was applied to PCoA-projected Bray–Curtis distances (first 10 principal coordinates, capturing >60% of variance). Optimal k was selected by maximum silhouette score over k = 2 to 5. Cluster characterisation used mean family-level composition within each cluster.

**S.M.6. Mantel Test**

Spearman rank correlations between geographic distance matrices (Haversine) and compositional distance matrices (Bray–Curtis) were computed using the Mantel test with 999 permutations. The test was performed on (a) all 1,672 samples from 1,456 unique individuals and (b) LogMPIE-only subjects (n =

1,004), where 14 cities were sampled with identical protocols, isolating geographic signal from batch effects.

**S.M.7. IGHI Weight Sensitivity Analysis**
Six alternative weighting schemes were evaluated against the default (Diversity 0.30, Balance 0.25, Core 0.20, Typicality 0.25):

| Scheme | Diversity | Balance | Core | Typicality |
|---|---|---|---|---|
| Default | 0.30 | 0.25 | 0.20 | 0.25 |
| Equal | 0.25 | 0.25 | 0.25 | 0.25 |
| Diversity-heavy | 0.50 | 0.20 | 0.15 | 0.15 |
| Balance-heavy | 0.15 | 0.40 | 0.20 | 0.25 |
| Core-heavy | 0.20 | 0.20 | 0.40 | 0.20 |
| Typicality-heavy | 0.20 | 0.20 | 0.15 | 0.45 |

For each scheme, IGHI was recomputed for all 1,672 samples from 1,456 unique individuals and Spearman rank correlation computed against the default ranking.

**S.M.8. Shotgun Metagenomic (WGS) Processing of the D'Aloisio Cohort**

**S.M.8.1. Data acquisition and computational environment**

Paired-end whole-genome shotgun (WGS) reads for the 110 samples of the D'Aloisio et al. cohort [16] (BioProject PRJNA1082632) were retrieved from the NCBI Sequence Read Archive using the SRA Toolkit (v3.4.1) and stored as gzip-compressed, per-accession FASTQ files. All analytical stages were run in isolated, version-pinned Conda (Miniforge3) environments: KneadData v0.10.0 with FastQC v0.12.1, Trimmomatic v0.41, Bowtie2 v2.5.5, SAMtools v1.24, SeqKit v2.13.0 and Seqtk v1.5 (environment *kneaddata_v010*); MetaPhlAn v4.2.5 and HUMAnN v3.9 (environment *metaphlan_env*, Python v3.10.20); and MultiQC v1.35 (environment *multiqc_env*, Python v3.14.7). Processing was performed on a single shared-memory Linux host (6 cores, 60 GB RAM) by batch scripts that terminate on unhandled errors during preprocessing, isolate per-sample failures during profiling, and resume safely after interruption. The complete software inventory is given in Table 1 of the Bioinformatics Software and Computational Environment record.

**S.M.8.2. Empirical identification of the sequencing adapter**

The source study states that adapters, primers and host reads were removed with KneadData, but the adapter reference, library-preparation chemistry and sequencing kit are reported neither in the article nor in its repository or supplementary materials. Because adapter identity is a required input to Trimmomatic's ILLUMINACLIP step, we established it empirically rather than assuming the Nextera adapter set bundled with KneadData, which an initial configuration had used by default. Verification was performed on a six-sample pilot (SRR28331611, SRR28331633, SRR28331652, SRR28331677, SRR28331696 and SRR28331728; 5.5% of the batch) in four phases.

*Phase 1.* KneadData was run twice per pilot sample, once without adapter clipping and once with the Nextera adapter file. Reads that survived as properly paired without clipping but were demoted or discarded under Nextera clipping were isolated by set difference of sorted read identifiers, yielding 68,311 reads whose fate depended on the Nextera assumption.

*Phase 2.* Each of these reads was tested exhaustively, with Trimmomatic's default 2-mismatch tolerance, against six Nextera adapter and orientation variants (19–33 bp) and the full-length TruSeq3 R1 and R2 adapters (34 bp). Neither returned a match (0/68,311 for every variant). A 13-bp generic Illumina universal-adapter prefix (AGATCGGAAGAGC) gave a marginal signal (0.24%; 0.26% on the full raw reads), roughly twice the random-match expectation for a probe of that length. It was not corroborated by Phase 3 and was excluded, since probes this short have an intrinsically high false-positive rate under Trimmomatic's seed-and-extend model.

*Phase 3.* AdapterRemoval3 (v3.0.1), run in report-only mode on each pilot sample, reconstructed adapter consensus sequences directly from paired-read overlaps without any prior hypothesis. The same consensus was recovered independently in all six samples (R1, 5′-…GGAATTCTCGGGTGCCAAGGAAC-3′; R2, 5′-…GATCGTCGGACT-3′), with a drop in per-base consensus quality at the adapter boundary, the expected signature of adapter read-through. The tool's internal reference matching labelled the R1 consensus as a small-RNA-library adapter. Because several small-RNA library kits share this adapter architecture, no single-kit attribution is claimed.

*Phase 4.* The consensus sequences were searched, with 2-mismatch tolerance, in the complete untrimmed read sets of all six samples. They were present in 2.0–3.2% of R1 and 2.6–4.0% of R2 reads, consistently across samples of approximately 3–10 million read pairs and independently of sequencing depth. Under the same framework the Nextera and TruSeq3 sequences were absent.

The adapters adopted for ILLUMINACLIP were therefore R1 5′-TGGAATTCTCGGGTGCCAAGGAAC-3′ (24 nt) and R2 5′-AGTCCGACGATCGTCGGACT-3′ (20 nt), applied with the originally intended stringency (2:30:10). In the full production pipeline, this configuration removed 1.02–1.66% of read pairs (mean ≈1.30%) relative to unclipped processing, again consistent across samples, whereas Nextera clipping affected only about 0.2% of pairs, in line with false-positive trimming. The selection deliberately favoured sensitivity, because uncorrected adapter contamination would propagate systematically into every downstream profile, whereas the cost of a false-positive clip is bounded and absorbed by the minimum-length filter.

Two limitations apply. First, the commercial kit of origin could not be determined from sequence alone. Low-level adapter carry-over from small-RNA libraries multiplexed on a shared flow cell is a plausible but unverified explanation. Second, verification was performed on six samples; their consistency supports, but does not prove, uniform applicability to all 110 samples.

##### S.M.8.3. Quality filtering and host-read depletion

Reads were processed with KneadData v0.10.0, which invokes Trimmomatic [32]. Trimming steps were executed in the following order: ILLUMINACLIP (confirmed adapters, 2:30:10), SLIDINGWINDOW:4:25, HEADCROP:10, CROP:120 and MINLEN:60. The last four parameters follow the source study. Host-derived reads were depleted by Bowtie2 v2.5.5 alignment [33] to a human reference index (hg_39) with --decontaminate-pairs lenient, which retains a pair unless both mates are host-derived. The tandem-repeat filtering step was bypassed, and FastQC was run on the final reads. A sample was accepted only if both paired output files were present. The retained files were compressed, and R1/R2 read counts recomputed from the compressed files were required to match and be non-zero. Per-sample results were aggregated with MultiQC [36].

##### S.M.8.4. Taxonomic profiling

Taxonomic profiles were generated with MetaPhlAn v4.2.5 [34] from the host-depleted paired reads, against the explicitly pinned marker database mpa_vJun23_CHOCOPhlAnSGB_202403 (Bowtie2 --bt2_ps sensitive, 2 threads per sample). The read-to-marker mapping output was retained for re-profiling. The retrieved FASTQ files carried a trailing comment on the "+" separator line that did not match the "@" identifier line, which caused MetaPhlAn's FASTQ parser to abort. For each sample, a working copy was therefore generated in which the separator line was replaced by a bare "+". The identifier on this line is optional under the FASTQ specification, so the change affects no sequence, quality or read-identity information. It was validated on a pilot sample (identical line counts; MetaPhlAn exit status 0). The upstream KneadData output was opened read-only and never modified. Samples were processed sequentially, because parallel execution caused memory-related terminations whose cause was not isolated.

##### S.M.8.5. Functional profiling

Gene-family abundances, pathway abundances and pathway coverage were estimated with HUMAnN v3.9 [35]. Each sample's MetaPhlAn profile restricted the nucleotide search to the detected taxa (ChocoPhlAn v201901_v31). Reads unmapped at that tier were searched by translated search against the EC-filtered UniRef90 database (release 201901b). Database paths were passed explicitly on every invocation, and 2 threads were used per sample. Input reads were the same header-corrected, host-depleted reads as in S.M.1.4, concatenated into one file as HUMAnN requires. HUMAnN's startup check of the MetaPhlAn version failed because the MetaPhlAn binary prints an extra database-status line, so a wrapper that returns a single version line was placed ahead of the binary on PATH. It changes no analytical parameter or database. Peak memory in the pilot sample was approximately 10.4 GB, and execution was sequential.

##### S.M.8.6. Provenance and quality assurance

Each profiling stage maintained a per-sample ledger of status, runtime, peak memory and exit code. A sample counted as successful only if the tool exited with status 0 and the primary output (MetaPhlAn profile; HUMAnN pathway-abundance table) was non-empty. Failed samples were retained with diagnostics and re-run from the start. Samples whose MetaPhlAn profile was missing were excluded from HUMAnN and recorded as such.

#### Supplementary Figures

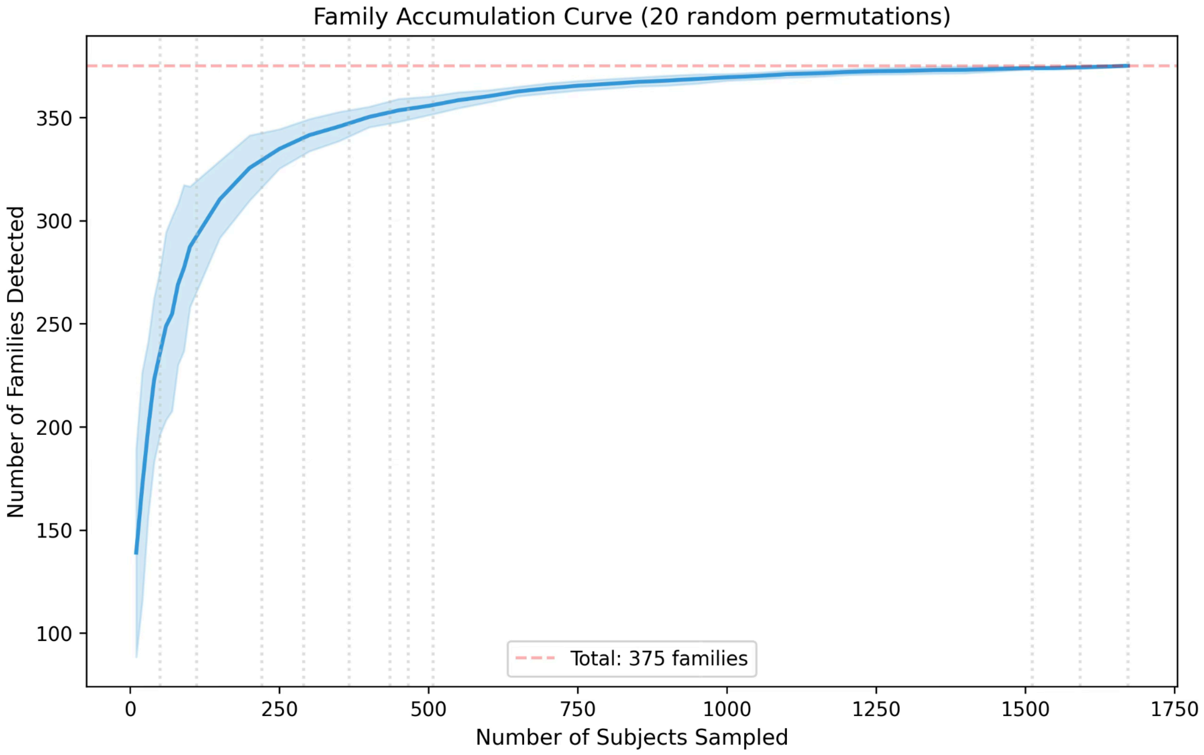


**Fig.** S1. Family accumulation (rarefaction) curve across all 1,672 subjects. The curve approaches saturation near 375 families, indicating adequate sampling of family-level diversity in the Indian healthy gut. Shaded area represents 95% confidence interval from 100 random subject orderings.

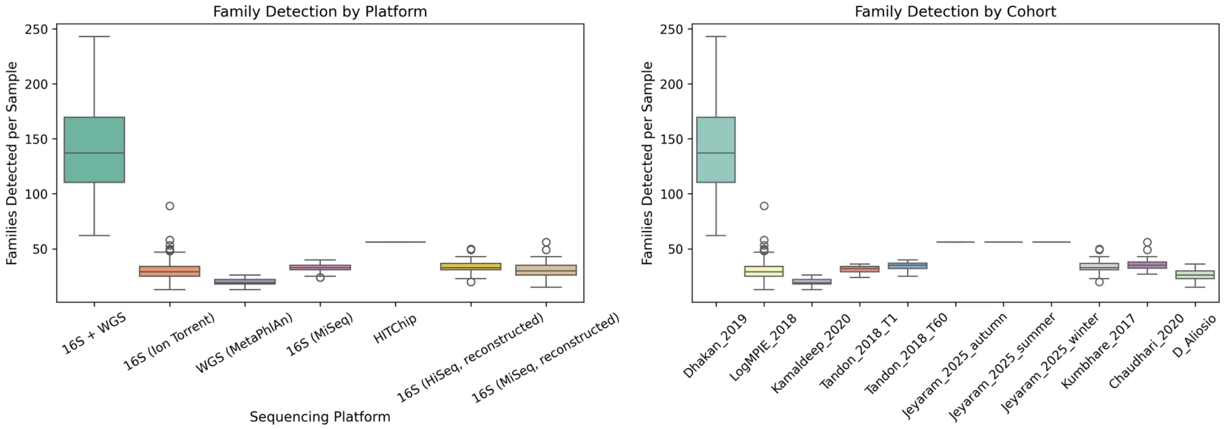


**Fig.** S2. Number of detected families per sample by sequencing platform (16S-Illumina, 16S-IonTorrent, HITChip, WGS). WGS samples (Dhakan cohort) detect substantially more families than 16S-based methods, explaining the high family richness variance observed in the Central zone (§3.2).

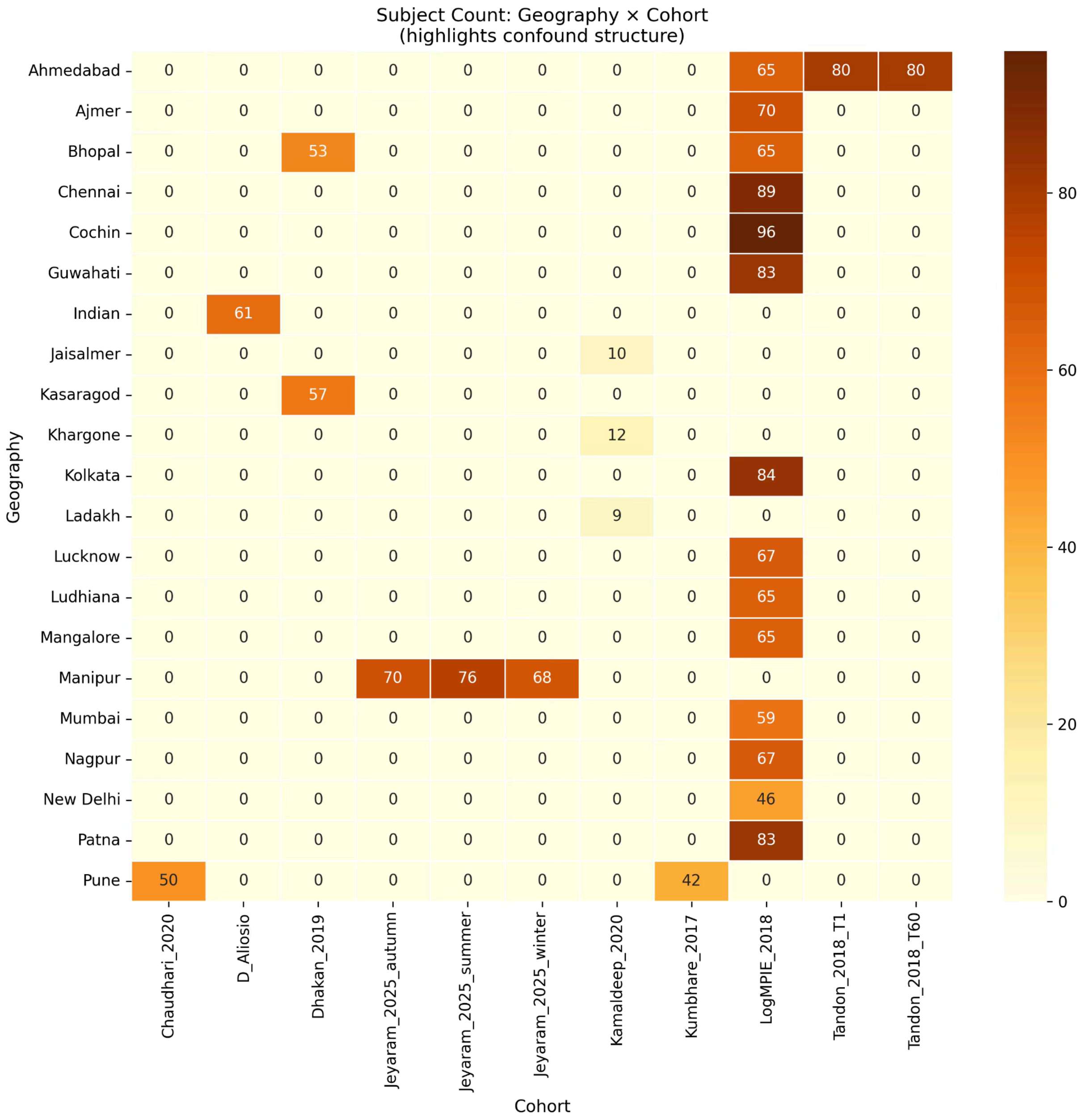


**Fig.** S3. Cohort × geography cross tabulation showing the number of subjects per cell. Twenty of 22 city-level geographies are sampled by a single cohort, illustrating the deeply confounded study–geography structure that drives the 30.4% shared variance in the partitioning analysis (Fig. 4).

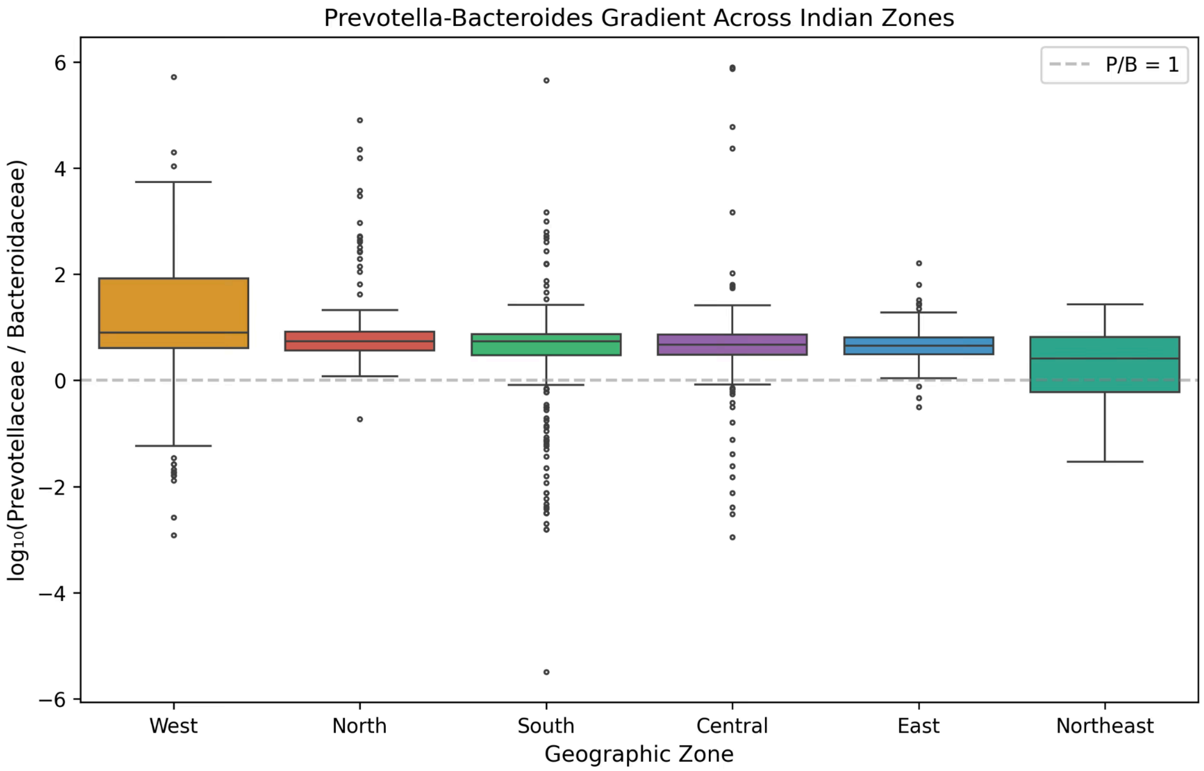


**Fig.** S4. $Log_{10}$(Prevotellaceae/Bacteroidaceae) ratio by geographic zone. Northeast subjects are overwhelmingly Prevotella-dominant; West subjects show the most balanced ratio. This gradient is consistent with the dietary continuum from fermented-food-rich diets of Manipur to diverse urban diets of western Indian cities.

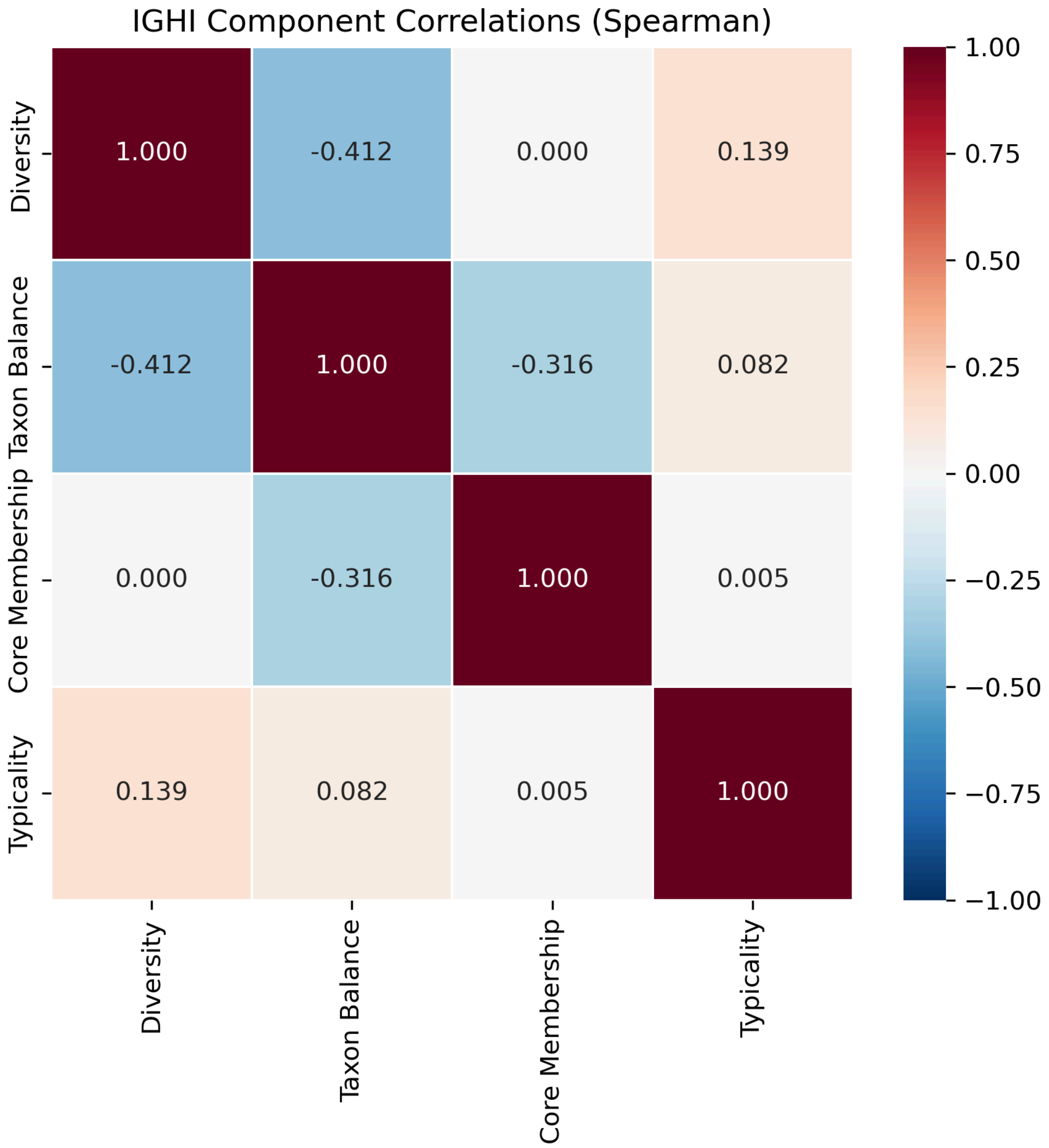


**Fig.** S5. *Pairwise Spearman correlations between the four IGHI components (Diversity, Taxon Balance, Core Membership, Community Typicality). Most pairs show weak correlation (|ρ| < 0.3), with the exception of Diversity vs. Taxon Balance (ρ = −0.41) and Taxon Balance vs. Core Membership (ρ = −0.32), confirming that the four components capture largely independent dimensions of gut health.*

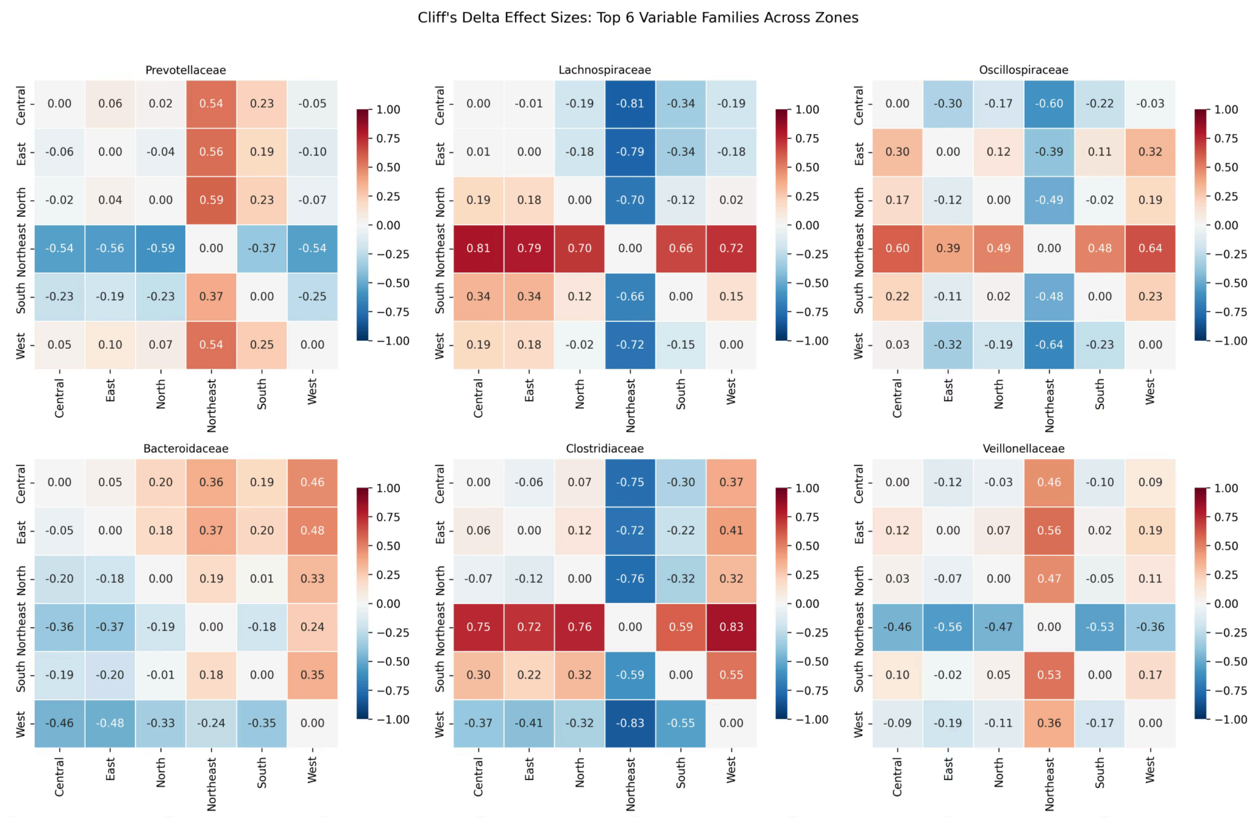


**Fig.** S6. Cliff's delta effect size matrix for the top 20 most variable families across all pairwise zone comparisons. Red indicates enrichment in the row zone relative to the column zone; blue indicates depletion. Of 300 pairwise comparisons, 216 (72%) were significant after FDR correction, with 56 (19%) showing large effect sizes (|δ| > 0.474).

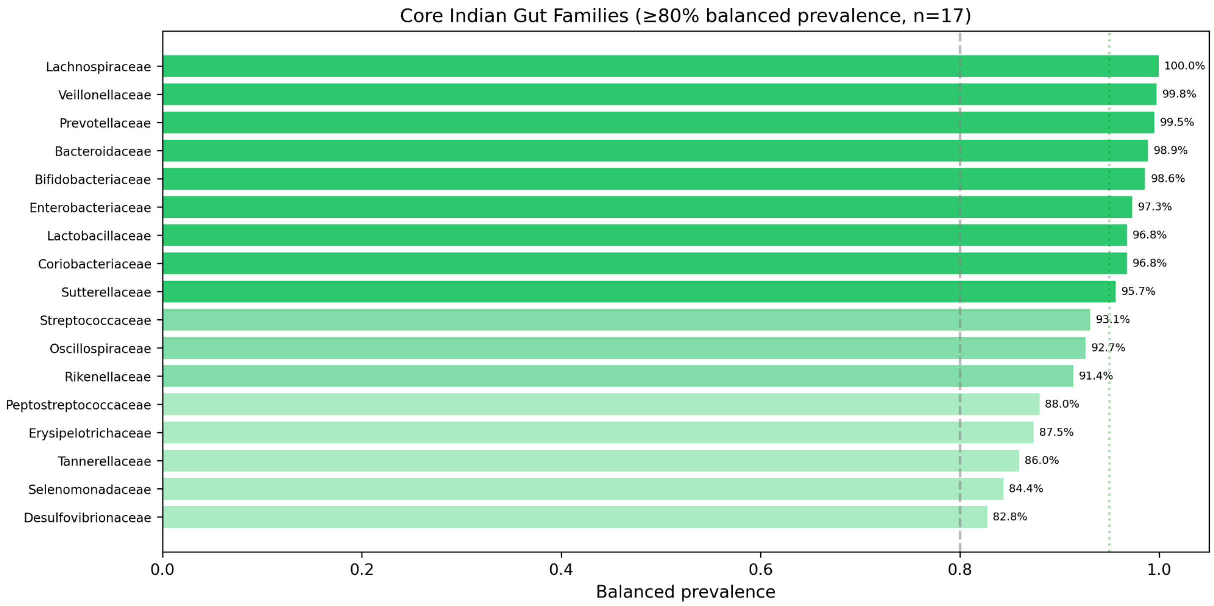


**Fig.** S7. The 17 core families of the healthy Indian gut, defined as those present in ≥80% balanced prevalence across cohorts. Bars show balanced prevalence (%). Lachnospiraceae reaches 100% prevalence; Desulfovibrionaceae at 82.8% is the threshold family.

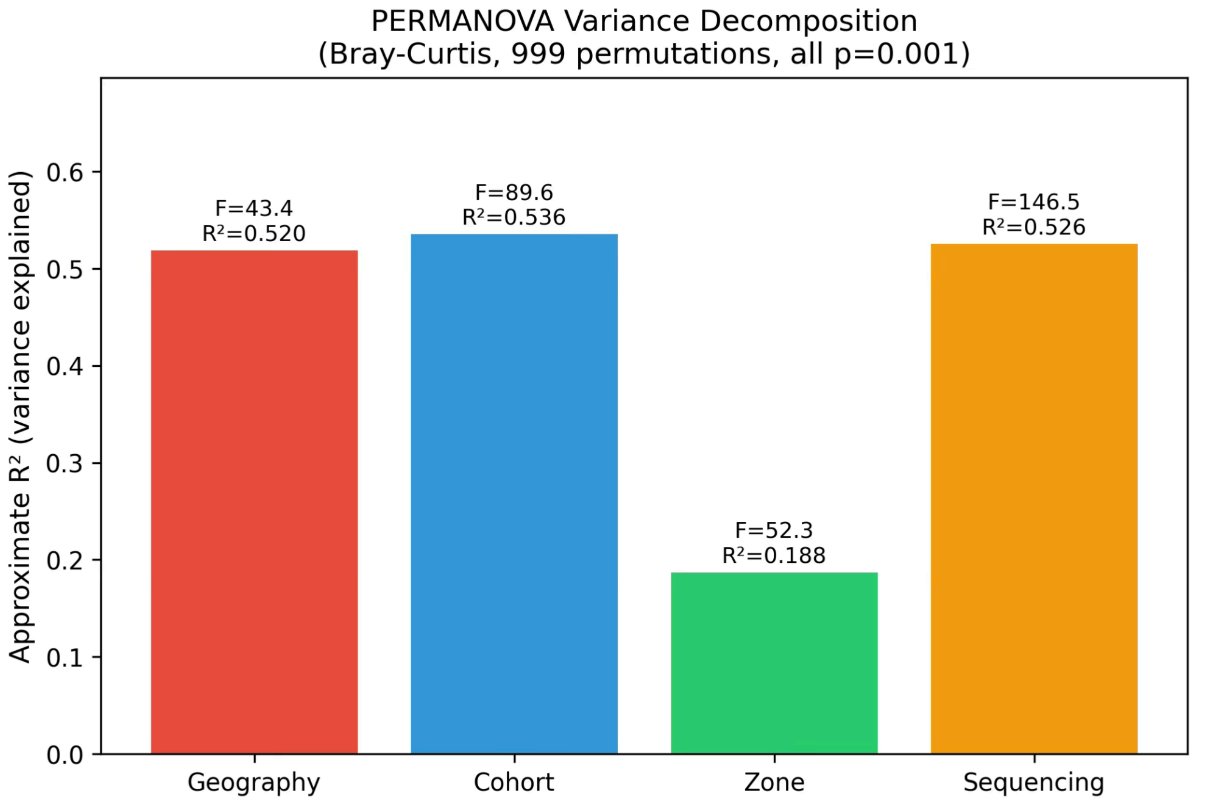


**Fig.** S8. PERMANOVA $R^2$ values for each grouping variable (geography, cohort, geographic zone, sequencing platform) computed on Bray–Curtis distances with 999 permutations. All tests significant ($p = 0.001$). Cohort and geography explain comparable variance, highlighting the confounded design.

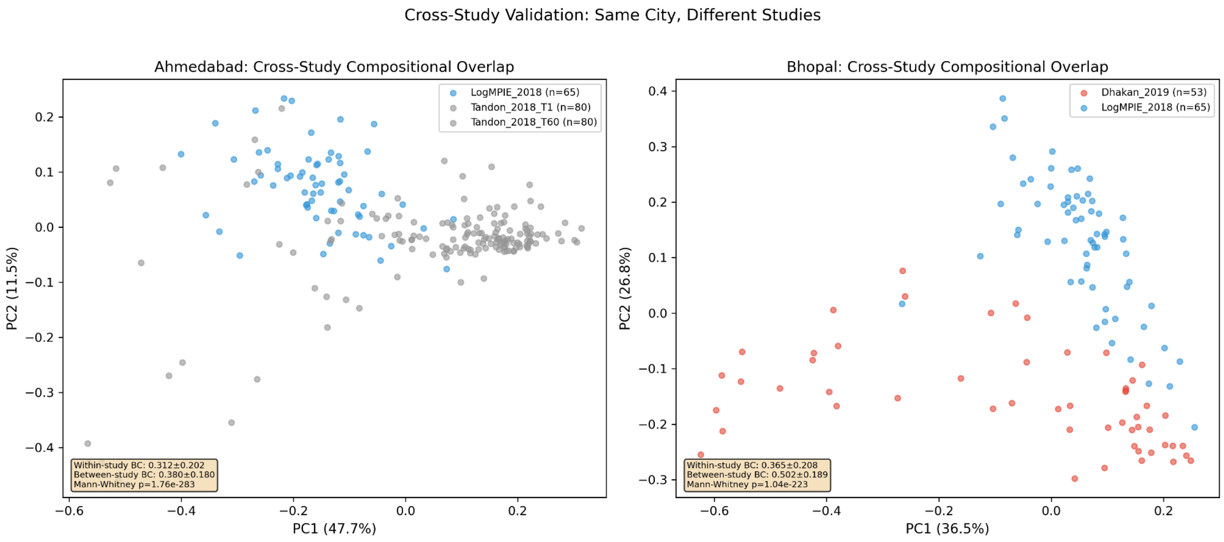

**Fig.** S9. PCoA comparison of subjects from geographies sampled by multiple cohorts: (a) Ahmedabad (Tandon + LogMPIE) and (b) Pune (Chaudhari + Kumbhare + LogMPIE). Points coloured by cohort. Overlapping distributions indicate genuine cross-study reproducibility at shared geographies; separation indicates residual batch effects.

### Supplementary Tables

| **Family** | **Prevalence (%)** | **Mean Rel. Abundance** | **Cohorts Detected** | **Core (≥80%)** |
|---|---|---|---|---|
| Lachnospiraceae | 100 | 0.0785 | 11 | Yes |
| Prevotellaceae | 99.76 | 0.3945 | 11 | Yes |
| Bacteroidaceae | 99.7 | 0.091 | 11 | Yes |
| Veillonellaceae | 99.52 | 0.0437 | 11 | Yes |
| Sutterellaceae | 98.68 | 0.0261 | 11 | Yes |
| Lactobacillaceae | 98.68 | 0.0208 | 11 | Yes |
| Oscillospiraceae | 98.5 | 0.1667 | 11 | Yes |
| Bifidobacteriaceae | 95.28 | 0.0168 | 11 | Yes |
| Enterobacteriaceae | 95.28 | 0.0131 | 11 | Yes |
| Rikenellaceae | 93.66 | 0.009 | 11 | Yes |
| Selenomonadaceae | 92.34 | 0.015 | 10 | Yes |
| Coriobacteriaceae | 90.55 | 0.0036 | 11 | Yes |
| Desulfovibrionaceae | 88.16 | 0.0028 | 11 | Yes |

| | | | | |
|---|---|---|---|---|
| Eubacteriaceae | 87.86 | 0.0272 | 9 | No |
| Tannerellaceae | 87.32 | 0.006 | 10 | Yes |
| Coprobacillaceae | 84.93 | 0.0033 | 10 | No |
| Pasteurellaceae | 84.69 | 0.0242 | 9 | No |
| Clostridiaceae | 82.78 | 0.0171 | 9 | No |
| Peptostreptococcaceae | 78.41 | 0.0015 | 11 | Yes |
| Acidaminococcaceae | 73.86 | 0.0038 | 9 | No |
| Actinomycetaceae | 71.23 | 2.20E-04 | 11 | No |
| Succinivibrionaceae | 69.44 | 0.0067 | 10 | No |
| Erysipelotrichaceae | 69.02 | 0.0011 | 11 | Yes |
| Streptococcaceae | 64.71 | 0.0068 | 11 | Yes |
| Akkermansiaceae | 63.94 | 0.0036 | 9 | No |
| Odoribacteraceae | 56.52 | 0.0011 | 7 | No |
| Turicibacteraceae | 55.2 | 6.90E-04 | 7 | No |
| Eggerthellaceae | 51.67 | 4.74E-04 | 9 | No |
| Treponemataceae | 51.08 | 0.0027 | 5 | No |
| Pseudomonadaceae | 50.24 | 6.02E-04 | 8 | No |
| Atopobiaceae | 46.35 | 7.97E-04 | 10 | No |

| | | | | |
|---|---|---|---|---|
| Campylobacteraceae | 45.93 | 2.10E-04 | 10 | No |
| Micrococcaceae | 45.39 | 1.29E-04 | 11 | No |
| Moraxellaceae | 43.66 | 1.98E-04 | 10 | No |
| Barnesiellaceae | 42.58 | 8.14E-04 | 7 | No |
| Fusobacteriaceae | 37.8 | 5.02E-04 | 7 | No |
| Carnobacteriaceae | 34.81 | 5.57E-05 | 9 | No |
| Oxalobacteraceae | 31.04 | 1.54E-04 | 8 | No |
| Gemellaceae | 29.07 | 2.94E-05 | 10 | No |
| Corynebacteriaceae | 27.81 | 6.88E-05 | 8 | No |
| Peptoniphilaceae | 26.97 | 1.33E-04 | 5 | No |
| Burkholderiaceae | 26.26 | 4.61E-05 | 8 | No |
| Butyricicoccaceae | 25.54 | 3.84E-04 | 7 | No |
| Aerococcaceae | 24.34 | 5.08E-05 | 7 | No |
| Helicobacteraceae | 24.04 | 1.19E-04 | 8 | No |
| Yersiniaceae | 23.8 | 9.12E-05 | 5 | No |
| Methylobacteriaceae | 23.74 | 1.38E-05 | 6 | No |
| Anoxybacillaceae | 22.01 | 1.95E-04 | 2 | No |
| Bacillaceae | 22.01 | 6.98E-05 | 6 | No |
| Vibrionaceae | 22.01 | 9.07E-05 | 5 | No |

*[Showing top 50 of 380 rows. Full table available in the GitHub repository.]*

**Table S1. NCBI Family-Level Taxonomy Mapping Table**
First 50 rows mapping of all input taxonomic labels from each cohort to harmonised NCBI family names. Columns: input label, source cohort, resolution step (exact match / synonym / genus-to-family fallback / Unassigned), matched NCBI family, NCBI TaxID, and any manual overrides.

The full table (all rows) is deposited with the dataset at:
https://github.com/TusharParihar128/BioMeBar_Indian_Healthy_Cohorts_datasets

| Family | Prevalence_pct | Mean_RelAbundance | Median_RelAbundance | SD_RelAbundance | Prevalence_Central_pct | Prevalence_East_pct | Prevalence_Indian_Unspecified_pct | Prevalence_North_pct | Prevalence_Northeast_pct | Prevalence_South_pct | Prevalence_West_pct |
|---|---|---|---|---|---|---|---|---|---|---|---|
| Lachnospiraceae | 100.00 | 0.0785 | 0.0482 | 0.0882 | 100.00 | 100.00 | 100.00 | 100.00 | 100.00 | 100.00 | 100.00 |
| Prevotellaceae | 99.76 | 0.3945 | 0.4126 | 0.2266 | 100.00 | 100.00 | 95.08 | 100.00 | 100.00 | 99.67 | 100.00 |
| Bacteroidaceae | 99.70 | 0.0910 | 0.0688 | 0.1010 | 98.98 | 100.00 | 100.00 | 99.50 | 100.00 | 99.67 | 99.77 |
| Veillonellaceae | 99.52 | 0.0437 | 0.0184 | 0.0779 | 99.49 | 100.00 | 98.36 | 99.00 | 100.00 | 100.00 | 99.10 |
| Sutterellaceae | 98.68 | 0.0261 | 0.0111 | 0.0500 | 100.00 | 100.00 | 93.44 | 99.00 | 100.00 | 99.67 | 96.61 |
| Lactobacillaceae | 98.68 | 0.0208 | 0.0130 | 0.0338 | 99.49 | 100.00 | 93.44 | 100.00 | 100.00 | 98.70 | 97.07 |
| Oscillospiraceae | 98.50 | 0.1667 | 0.1513 | 0.1109 | 95.94 | 100.00 | 100.00 | 91.50 | 100.00 | 100.00 | 100.00 |

| | | | | | | | | | | | |
|---|---|---|---|---|---|---|---|---|---|---|---|
| Enterobacteriaceae | 95.28 | 0.0131 | 0.0027 | 0.0487 | 94.92 | 94.01 | 93.44 | 88.50 | 100.00 | 93.49 | 97.29 |
| Bifidobacteriaceae | 95.28 | 0.0168 | 0.0029 | 0.0487 | 94.92 | 91.02 | 98.36 | 88.00 | 100.00 | 97.07 | 95.49 |
| Rikenellaceae | 93.66 | 0.0090 | 0.0041 | 0.0166 | 94.92 | 90.42 | 86.89 | 92.50 | 100.00 | 97.72 | 88.71 |
| Selenomonadaceae | 92.34 | 0.0150 | 0.0038 | 0.0364 | 90.86 | 98.20 | 78.69 | 84.00 | 99.66 | 96.09 | 88.94 |
| Coriobacteriaceae | 90.55 | 0.0036 | 0.0012 | 0.0095 | 91.88 | 80.24 | 95.08 | 85.50 | 100.00 | 92.18 | 88.04 |
| Desulfovibrionaceae | 88.16 | 0.0028 | 7.15e-04 | 0.0073 | 91.88 | 93.41 | 65.57 | 87.00 | 99.66 | 94.79 | 75.85 |
| Eubacteriaceae | 87.86 | 0.0272 | 0.0181 | 0.0305 | 100.00 | 100.00 | 98.36 | 100.00 | 100.00 | 99.67 | 54.63 |
| Tannerellaceae | 87.32 | 0.0060 | 0.0018 | 0.0116 | 81.22 | 82.04 | 81.97 | 73.00 | 99.66 | 91.86 | 87.81 |
| Coprobacillaceae | 84.93 | 0.0033 | 0.0013 | 0.0069 | 87.31 | 86.83 | 54.10 | 75.50 | 99.66 | 85.99 | 81.04 |
| Pasteurellaceae | 84.69 | 0.0242 | 0.0013 | 0.0585 | 89.34 | 90.42 | 0.0000 | 86.50 | 100.00 | 92.83 | 75.40 |

| | | | | | | | | | | | |
|---|---|---|---|---|---|---|---|---|---|---|---|
| Clostridiaceae | 82.78 | 0.0171 | 0.0052 | 0.0315 | 94.92 | 97.01 | 9.84 | 90.50 | 100.00 | 99.35 | 55.53 |
| Peptostreptococcaceae | 78.41 | 0.0015 | 1.93e-04 | 0.0106 | 81.73 | 68.86 | 90.16 | 60.50 | 96.97 | 80.13 | 73.36 |
| Acidaminococcaceae | 73.86 | 0.0038 | 4.40e-04 | 0.0160 | 81.73 | 81.44 | 85.25 | 70.00 | 98.65 | 85.02 | 43.34 |
| Actinomycetaceae | 71.23 | 2.20e-04 | 4.02e-05 | 6.42e-04 | 75.63 | 59.88 | 26.23 | 50.50 | 95.62 | 75.57 | 69.75 |
| Succinivibrionaceae | 69.44 | 0.0067 | 1.00e-04 | 0.0338 | 60.41 | 76.05 | 6.56 | 57.00 | 98.32 | 59.93 | 72.46 |
| Erysipelotrichaceae | 69.02 | 0.0011 | 1.96e-04 | 0.0057 | 68.02 | 50.30 | 75.41 | 53.50 | 94.95 | 68.08 | 65.91 |
| Streptococcaceae | 64.71 | 0.0068 | 3.50e-04 | 0.0252 | 54.31 | 19.76 | 88.52 | 23.00 | 98.99 | 67.10 | 77.20 |
| Akkermansiaceae | 63.94 | 0.0036 | 1.66e-04 | 0.0305 | 63.45 | 55.69 | 65.57 | 56.00 | 98.65 | 75.24 | 39.50 |
| Odoribacteraceae | 56.52 | 0.0011 | 8.71e-05 | 0.0028 | 68.53 | 44.91 | 68.85 | 53.50 | 23.91 | 74.59 | 64.56 |
| Turicibacteraceae | 55.20 | 6.90e-04 | 2.67e-05 | 0.0029 | 70.56 | 58.68 | 31.15 | 47.00 | 22.90 | 64.82 | 69.07 |

| | | | | | | | | | | | |
|---|---|---|---|---|---|---|---|---|---|---|---|
| Eggerthellaceae | 51.67 | 4.74e-04 | 1.93e-05 | 0.0031 | 58.88 | 37.13 | 62.30 | 28.50 | 83.50 | 50.16 | 42.66 |
| Treponemataceae | 51.08 | 0.0027 | 1.14e-05 | 0.0154 | 76.65 | 94.61 | 1.64 | 59.50 | 27.27 | 78.83 | 23.02 |
| Pseudomonadaceae | 50.24 | 6.02e-04 | 3.49e-06 | 0.0060 | 53.30 | 18.56 | 0.0000 | 23.50 | 87.54 | 69.06 | 41.76 |
| Atopobiaceae | 46.35 | 7.97e-04 | 0.0000 | 0.0086 | 49.24 | 31.14 | 27.87 | 15.00 | 79.80 | 38.11 | 50.79 |
| Campylobacteraceae | 45.93 | 2.10e-04 | 0.0000 | 0.0012 | 56.85 | 23.95 | 8.20 | 22.00 | 81.14 | 48.86 | 39.73 |
| Micrococcaceae | 45.39 | 1.29e-04 | 0.0000 | 0.0012 | 50.76 | 17.37 | 4.92 | 21.50 | 80.81 | 38.76 | 50.79 |
| Moraxellaceae | 43.66 | 1.98e-04 | 0.0000 | 0.0024 | 49.75 | 28.74 | 3.28 | 28.50 | 78.79 | 42.35 | 36.34 |
| Barnesiellaceae | 42.58 | 8.14e-04 | 0.0000 | 0.0030 | 46.19 | 31.14 | 39.34 | 36.00 | 25.93 | 59.61 | 48.08 |
| Fusobacteriaceae | 37.80 | 5.02e-04 | 0.0000 | 0.0052 | 53.81 | 32.34 | 11.48 | 16.00 | 22.22 | 49.51 | 48.53 |
| Carnobacteriaceae | 34.81 | 5.57e-05 | 0.0000 | 0.0011 | 31.98 | 8.38 | 0.0000 | 8.00 | 81.14 | 28.99 | 35.89 |

| | | | | | | | | | | | |
|---|---|---|---|---|---|---|---|---|---|---|---|
| Oxalobacteraceae | 31.04 | 1.54e-04 | 0.0000 | 7.02e-04 | 39.09 | 11.98 | 0.0000 | 10.00 | 84.18 | 36.48 | 9.03 |
| Gemellaceae | 29.07 | 2.94e-05 | 0.0000 | 1.65e-04 | 26.40 | 0.5988 | 8.20 | 1.00 | 72.05 | 18.57 | 34.99 |
| Corynebacteriaceae | 27.81 | 6.88e-05 | 0.0000 | 8.23e-04 | 38.58 | 10.18 | 0.0000 | 12.50 | 74.75 | 28.01 | 8.80 |
| Peptoniphilaceae | 26.97 | 1.33e-04 | 0.0000 | 8.46e-04 | 50.76 | 26.35 | 13.11 | 20.50 | 14.48 | 46.91 | 16.03 |
| Burkholderiaceae | 26.26 | 4.61e-05 | 0.0000 | 4.12e-04 | 31.47 | 4.19 | 0.0000 | 5.50 | 74.07 | 23.13 | 15.35 |
| Butyricicoccaceae | 25.54 | 3.84e-04 | 0.0000 | 0.0018 | 29.44 | 2.99 | 95.08 | 0.0000 | 0.0000 | 19.22 | 55.76 |
| Aerococcaceae | 24.34 | 5.08e-05 | 0.0000 | 6.65e-04 | 31.98 | 6.59 | 0.0000 | 10.50 | 73.74 | 22.80 | 5.19 |
| Helicobacteraceae | 24.04 | 1.19e-04 | 0.0000 | 0.0010 | 25.38 | 3.59 | 1.64 | 6.00 | 73.40 | 14.98 | 15.58 |
| Yersiniaceae | 23.80 | 9.12e-05 | 0.0000 | 7.99e-04 | 36.04 | 1.80 | 0.0000 | 3.50 | 73.40 | 28.34 | 2.71 |
| Methylobacteriaceae | 23.74 | 1.38e-05 | 0.0000 | 1.29e-04 | 23.35 | 1.80 | 0.0000 | 4.00 | 73.06 | 15.64 | 16.93 |

| | | | | | | | | | | | |
|---|---|---|---|---|---|---|---|---|---|---|---|
| Vibrionaceae | 22.01 | 9.07e-05 | 0.0000 | 0.0012 | 32.99 | 1.20 | 0.0000 | 6.50 | 75.76 | 18.57 | 1.35 |
| Bacillaceae | 22.01 | 6.98e-05 | 0.0000 | 7.97e-04 | 27.92 | 4.79 | 0.0000 | 1.50 | 75.42 | 22.15 | 2.26 |
| Anoxybacillaceae | 22.01 | 1.95e-04 | 0.0000 | 9.32e-04 | 37.56 | 2.99 | 0.0000 | 6.50 | 25.59 | 52.12 | 9.03 |

*[Showing top 50 of 380 rows. Full table available in the GitHub repository.]*

**Table S2. Complete Core Family Statistics**
All 380 families with prevalence (%), mean relative abundance, median relative abundance, standard deviation, and per-zone prevalence values. Available in the GitHub repository.

| Test | pseudo_F | p_value | permutations | method |
|---|---|---|---|---|
| Geography (standard) | 43.4095 | 0.0010 | 999 | unrestricted |
| Cohort (standard) | 89.5561 | 0.0010 | 999 | unrestricted |
| Zone (standard) | 52.2909 | 0.0010 | 999 | unrestricted |
| Geography \| Cohort (restricted) | 43.4095 | 0.0010 | 999 | within-cohort |
| Zone \| Cohort (restricted) | 52.2909 | 0.8960 | 999 | within-cohort |

**Table S3. Partial PERMANOVA Results**
Restricted permutation PERMANOVA results testing geographic signal after controlling for cohort effects.

| Factor | KW_statistic | p_value | n_groups | min_dispersion_group | max_dispersion_group |
|---|---|---|---|---|---|
| Geography | 515.5525 | 1.61e-96 | 21 | Guwahati (0.146) | Pune (0.390) |

| | | | | | |
|---|---|---|---|---|---|
| Cohort | 344.2608 | 6.58e-68 | 11 | Tandon_2018_T60 (0.179) | Kumbhare_2017 (0.380) |
| Zone | 156.2423 | 3.70e-31 | 7 | East (0.223) | Indian_Unspecified (0.375) |

**Table S4. PERMDISP Results**
Multivariate dispersion heterogeneity test results. Significant PERMDISP indicates that PERMANOVA results may partly reflect dispersion differences rather than location shifts.

| **Family** | **Indicator_Zone** | **IndVal** | **Specificity** | **Fidelity** | **p_value** | **q_value** | **significant** |
|---|---|---|---|---|---|---|---|
| Streptococcaceae | Northeast | 0.7476 | 0.7552 | 0.9899 | 0.0010 | 0.0016 | True |
| Budviciaceae | Northeast | 0.7205 | 1.0000 | 0.7205 | 0.0010 | 0.0016 | True |
| Oceanospirillaceae | Northeast | 0.7138 | 0.9906 | 0.7205 | 0.0010 | 0.0016 | True |
| Peptococcaceae | Northeast | 0.7120 | 0.9881 | 0.7205 | 0.0010 | 0.0016 | True |
| Anaerovoracaceae | Northeast | 0.7108 | 0.9865 | 0.7205 | 0.0010 | 0.0016 | True |
| Paenibacillaceae | Northeast | 0.6975 | 0.9635 | 0.7239 | 0.0010 | 0.0016 | True |
| Enterococcaceae | Northeast | 0.6478 | 0.8948 | 0.7239 | 0.0010 | 0.0016 | True |
| Brachyspiraceae | Northeast | 0.6290 | 0.8730 | 0.7205 | 0.0010 | 0.0016 | True |
| Bacillaceae | Northeast | 0.6213 | 0.8238 | 0.7542 | 0.0010 | 0.0016 | True |
| Propionibacteriaceae | Northeast | 0.6028 | 0.8366 | 0.7205 | 0.0010 | 0.0016 | True |
| Aeromonadaceae | Northeast | 0.5919 | 0.8177 | 0.7239 | 0.0010 | 0.0016 | True |

| | | | | | | | |
|---|---|---|---|---|---|---|---|
| Carnobacteriaceae | Northeast | 0.5406 | 0.6662 | 0.8114 | 0.0010 | 0.0016 | True |
| Tannerellaceae | Northeast | 0.5226 | 0.5244 | 0.9966 | 0.0010 | 0.0016 | True |
| Oxalobacteraceae | Northeast | 0.5085 | 0.6041 | 0.8418 | 0.0010 | 0.0016 | True |
| Butyricicoccaceae | West | 0.5085 | 0.9121 | 0.5576 | 0.0010 | 0.0016 | True |
| Succinivibrionaceae | West | 0.4933 | 0.6808 | 0.7246 | 0.0010 | 0.0016 | True |
| Clostridiaceae | Northeast | 0.4817 | 0.4817 | 1.0000 | 0.0010 | 0.0016 | True |
| Staphylococcaceae | Northeast | 0.4804 | 0.6667 | 0.7205 | 0.0010 | 0.0016 | True |
| Treponemataceae | East | 0.4292 | 0.4536 | 0.9461 | 0.0010 | 0.0016 | True |
| Gemellaceae | Northeast | 0.4147 | 0.5756 | 0.7205 | 0.0010 | 0.0016 | True |
| Morganellaceae | Northeast | 0.4132 | 0.5504 | 0.7508 | 0.0010 | 0.0016 | True |
| Atopobiaceae | West | 0.4122 | 0.8116 | 0.5079 | 0.0010 | 0.0016 | True |
| Peptostreptococcaceae | West | 0.4108 | 0.5599 | 0.7336 | 0.0010 | 0.0016 | True |
| Lachnospiraceae | Northeast | 0.4082 | 0.4082 | 1.0000 | 0.0010 | 0.0016 | True |
| Desulfovibrionaceae | East | 0.3623 | 0.3878 | 0.9341 | 0.0010 | 0.0016 | True |
| Burkholderiaceae | Northeast | 0.3341 | 0.4511 | 0.7407 | 0.0010 | 0.0016 | True |
| Sphingomonadaceae | Northeast | 0.3235 | 0.4469 | 0.7239 | 0.0010 | 0.0016 | True |

| | | | | | | | |
|---|---|---|---|---|---|---|---|
| Aerococcaceae | Northeast | 0.3113 | 0.4222 | 0.7374 | 0.0010 | 0.0016 | True |
| Enterobacteriaceae | Central | 0.3088 | 0.3253 | 0.9492 | 0.0010 | 0.0016 | True |
| Pasteurellaceae | South | 0.2993 | 0.3224 | 0.9283 | 0.0010 | 0.0016 | True |

[Showing first 30 of 375 rows. Full table available in the GitHub repository.]

**Table S5. Indicator Species (IndVal) Results**
Top 30 indicator families by IndVal score (of 319 significant). Full table (375 rows) available in the repository.

| Test | n_samples | Spearman_r | p_value |
|---|---|---|---|
| All samples | 1611 | 0.1734 | 0.0010 |
| LogMPIE only | 1004 | 0.0087 | 0.4360 |

**Table S6. Mantel Test Results**
Geographic distance-decay analysis. The significant all-sample correlation disappears within the LogMPIE cohort (identical protocols across 14 cities), indicating the signal is batch-driven.

| Component_1 | Component_2 | Spearman_rho | p_value |
|---|---|---|---|
| Diversity | Taxon Balance | -0.4125 | 1.09E-69 |
| Diversity | Core Membership | 0.000262 | 0.991445 |
| Diversity | Typicality | 0.139099 | 1.12E-08 |
| Taxon Balance | Core Membership | -0.31635 | 3.55E-40 |
| Taxon Balance | Typicality | 0.081796 | 0.000815 |
| Core Membership | Typicality | 0.005148 | 0.833403 |

**Table S7. IGHI Component Correlation Matrix**
Pairwise Spearman correlations between the four IGHI components.

| Family | Zone_1 | Zone_2 | Cliffs_delta | Effect_size | MannWhitney_p | q_value | significant |
|---|---|---|---|---|---|---|---|
| Prevotellaceae | Central | East | 0.0630 | negligible | 0.3008 | 0.3498 | False |
| Prevotellaceae | Central | North | 0.0200 | negligible | 0.7300 | 0.7626 | False |
| Prevotellaceae | Central | Northeast | 0.5400 | large | 2.91e-24 | 1.78e-23 | True |
| Prevotellaceae | Central | South | 0.2250 | small | 1.93e-05 | 3.84e-05 | True |
| Prevotellaceae | Central | West | -0.0480 | negligible | 0.3289 | 0.3766 | False |
| Prevotellaceae | East | North | -0.0450 | negligible | 0.4542 | 0.5028 | False |
| Prevotellaceae | East | Northeast | 0.5620 | large | 8.65e-24 | 5.19e-23 | True |
| Prevotellaceae | East | South | 0.1880 | small | 7.38e-04 | 0.0012 | True |
| Prevotellaceae | East | West | -0.0980 | negligible | 0.0627 | 0.0817 | False |
| Prevotellaceae | North | Northeast | 0.5860 | large | 1.51e-28 | 1.13e-27 | True |
| Prevotellaceae | North | South | 0.2320 | small | 1.04e-05 | 2.12e-05 | True |
| Prevotellaceae | North | West | -0.0660 | negligible | 0.1769 | 0.2157 | False |
| Prevotellaceae | Northeast | South | -0.3740 | medium | 1.89e-15 | 7.79e-15 | True |
| Prevotellaceae | Northeast | West | -0.5420 | large | 7.68e-36 | 8.86e-35 | True |
| Prevotellaceae | South | West | -0.2470 | small | 8.63e-09 | 2.17e-08 | True |

| | | | | | | | |
|---|---|---|---|---|---|---|---|
| Lachnospiraceae | Central | East | -0.0060 | negligible | 0.9196 | 0.9415 | False |
| Lachnospiraceae | Central | North | -0.1920 | small | 9.11e-04 | 0.0015 | True |
| Lachnospiraceae | Central | Northeast | -0.8080 | large | 3.02e-52 | 1.01e-50 | True |
| Lachnospiraceae | Central | South | -0.3380 | medium | 1.42e-10 | 4.26e-10 | True |
| Lachnospiraceae | Central | West | -0.1910 | small | 1.15e-04 | 2.08e-04 | True |
| Lachnospiraceae | East | North | -0.1800 | small | 0.0029 | 0.0045 | True |
| Lachnospiraceae | East | Northeast | -0.7920 | large | 1.52e-45 | 3.04e-44 | True |
| Lachnospiraceae | East | South | -0.3350 | medium | 1.74e-09 | 4.65e-09 | True |
| Lachnospiraceae | East | West | -0.1840 | small | 4.62e-04 | 7.78e-04 | True |
| Lachnospiraceae | North | Northeast | -0.7040 | large | 1.84e-40 | 2.91e-39 | True |
| Lachnospiraceae | North | South | -0.1220 | negligible | 0.0202 | 0.0290 | True |
| Lachnospiraceae | North | West | 0.0170 | negligible | 0.7321 | 0.7626 | False |
| Lachnospiraceae | Northeast | South | 0.6570 | large | 2.44e-44 | 4.31e-43 | True |
| Lachnospiraceae | Northeast | West | 0.7190 | large | 6.76e-62 | 3.38e-60 | True |
| Lachnospiraceae | South | West | 0.1470 | small | 6.06e-04 | 0.0010 | True |

[Showing first 30 of 300 rows. Full table available in the GitHub repository.]

**Table S8. Consolidated Cliff's Delta Effect Sizes**

Pairwise Cliff's delta values for the top 20 most variable families across all zone pairs.

| Cluster | n | Dominant_Family | Top3_Families | Top_Zone | Top_Zone_Fraction |
|---|---|---|---|---|---|
| 1 | 519 | Oscillospiraceae | Oscillospiraceae (0.244); Lachnospiraceae (0.149); Prevotellaceae (0.126) | Northeast | 0.4046 |
| 2 | 1153 | Prevotellaceae | Prevotellaceae (0.515); Oscillospiraceae (0.132); Bacteroidaceae (0.079) | West | 0.3053 |

Silhouette scores for k = 2 to 6:

| k | silhouette |
|---|---|
| 2.0000 | 0.3434 |
| 3.0000 | 0.1978 |
| 4.0000 | 0.2212 |
| 5.0000 | 0.2328 |
| 6.0000 | 0.2018 |

**Table S9. Community Type (Enterotype) Assignments**
Two community types identified by k-medoids clustering (optimal k = 2).

| Unnamed: 0 | mean | std | median |
|---|---|---|---|
| Default (30/25/20/25) | 60.89501 | 10.35807 | 62.05043 |
| Equal (25/25/25/25) | 62.94213 | 9.590644 | 64.18978 |

| | | | |
|---|---|---|---|
| Diversity-heavy (50/20/15/15) | 57.84004 | 13.77674 | 58.80004 |
| Balance-heavy (15/40/20/25) | 60.86809 | 11.75926 | 62.3335 |
| Typicality-heavy (20/20/15/45) | 60.89048 | 10.95825 | 62.85988 |
| Core-heavy (20/20/40/20) | 68.58407 | 8.027474 | 69.55315 |

**Table S10. IGHI Weight Sensitivity Summary**
IGHI score distribution under six alternative weighting schemes.

**Data and Code Availability**
All data, scripts, and analysis outputs are available at:
https://github.com/TusharParihar128/BioMeBar_Indian_Healthy_Cohorts_datasets

| **File** | **Description** |
|---|---|
| data/BioMeBar_Unified_Family_Matrix.csv | Harmonised 1,672 × 387 matrix (7 metadata + 380 families) |
| data/BioMeBar_Unified_Metadata.csv | Subject metadata |
| scripts/06_publication_figures.py | Generates all main and supplementary figures |
| scripts/07_statistical_analyses.py | Runs all statistical analyses |
| results/PUBLICATION_FIGURES/ | Generated figure files (PNG + PDF) |
| results/STATISTICAL_ANALYSIS/ | Analysis output CSVs and figures |

### References


[16] L. D. D'Aloisio et al., "The adoption of a westernized gut microbiome in Indian Immigrants and Indo-Canadians is associated with dietary acculturation," *NPJ Biofilms Microbiomes*, vol. 11, no. 1, Dec. 2025, doi: 10.1038/s41522-025-00778-8

[28] M. Dufrêne and P. Legendre, "SPECIES ASSEMBLAGES AND INDICATOR SPECIES:THE NEED FOR A FLEXIBLE ASYMMETRICAL APPROACH," *Ecol. Monogr.*, vol. 67, no. 3, pp. 345–366, Aug. 1997, doi: https://doi.org/10.1890/0012-9615(1997)067[0345:SAAIST]2.0.CO;2.

[32] Bolger AM et al. Bioinformatics 30:2114–2120 (2014), doi:10.1093/bioinformatics/btu170

[33] Langmead B, Salzberg SL. Nat Methods 9:357–359 (2012), doi:10.1038/nmeth.1923

[34] Blanco-Míguez A et al. Nat Biotechnol 41:1633–1644 (2023), doi:10.1038/s41587-023-01688-w

[35] Beghini F et al. eLife 10:e65088 (2021), doi:10.7554/eLife.65088

[36] Ewels P et al. Bioinformatics 32:3047–3048 (2016), doi:10.1093/bioinformatics/btw354
